# Socioeconomic and Genomic Roots of Verbal Ability

**DOI:** 10.1101/544411

**Authors:** Guang Guo, Meng-Jung Lin, Kathleen M. Harris

## Abstract

Cognitive ability is one of the most potent and contentious human traits. Many issues surrounding cognitive ability especially those related to heredity is highly charged. Yet, all of the discussion on heredity has been based on non-DNA evidence. It is largely neglected that DNA and environmental data at individual level are indispensable for understanding the development of cognitive ability. In this article, we report findings from a study that uses both ability-related polygenic scores (PGSs) and a rich set of socioeconomic measures from Add Health. In an all-ethnicity sample excluding blacks, a social-science model predicts verbal ability well yielding an *R*^*2*^ of 17.5%. Adding two ability-related PGSs increases this *R*^*2*^ by 1.7%. Such models yield more accurate estimates of the effects of the PGSs and those of SES context, and provide an estimated degree to which SES context is influenced by parental genomes. Schooling and neighborhood remain important to verbal ability even after an early measure of verbal ability is adjusted in the model. Although the influence from the genome is evident, the influences of SES context are critical and cannot be dismissed.

## INTRODUCTION

Cognitive ability is one of the most potent human traits. It is potent for at least three reasons. Arguably, cognitive ability is the hallmark of humans that separates them from the other species in the natural world. It is ubiquitous and vital for basic subsistence in deep human history and for contemporary physical and economic well-being. Evidence suggests that more than an inconsequential portion of ability is related to heredity (Bouchard 1998; Bouchard and McGue 1981). Cognitive ability is as controversial as it is potent. Many issues surrounding cognitive ability especially those related to heredity is highly charged (e.g., Fischer et al. 1996; Herrnstein and Murray 1994; Jensen 1969). Yet, all of the empirical analysis concerning heredity has been based on non-DNA data. It is largely neglected that DNA evidence in combination with socioeconomic (SES) contextual information at individual level is indispensable for understanding how heredity and SES context jointly and interactively mold human cognitive ability. Such DNA evidence just became available recently (Lee et al. 2018; Okbay et al. 2016; Savage et al. 2018; Sniekers et al. 2017).

Understanding the roots of cognitive ability is relevant and important. Cognitive ability has been shown to be one of the best predictors of life outcomes such as educational attainment, occupation achievement, income, wealth, and health (e.g., Farkas et al. 1997; Farkas and Vicknair 1996; Jencks et al. 1979; Taubman and Wales 1974; Taubman and Wales 1972; Wraw et al. 2015). Cognitive ability remains important even with the evidence that its predictive power for later life outcomes tends to be markedly reduced once educational attainment is adjusted. This is so because cognitive ability is a key predictor of academic performance in school as well as educational attainment itself and because it exerts an effect on labor market outcomes independently from educational attainment (Jencks et al. 1979; Kerckhoff, Raudenbush and Glennie 2001; Rosenbaum 2001; Spilerman and Lunde 1991). Tests of cognitive ability and related cognitive achievement are routinely and nearly universally used in elementary and secondary education, college admissions, and admissions of graduate schools and professional schools in the United States (Hauser 2010; Lemann 1999).

Views differ enormously regarding what gives rise to cognitive ability. At the heart of the matter is how much weight should be placed on inheritance relative to after-birth learning. One long-standing and influential position considered it an “…inborn, all around intellectual ability … inherited, but not due to teaching or training.. uninfluenced by industry or zeal. (Burt et al. 1934, cited by RE Nisbett 2009).” Jensen maintained that “… the means for changing intelligence per se lie in the province of biology rather than psychology or education (1969).” Herrnstein and Murray (1994) expressed a similar assessment that the differences in intelligence were mostly at the hand of nature and there was little that government policies could change. As Nisbett (2009, p.36) pointed out, many prominent scholars believed that family environments have essentially nil effects, including JR Harris (1998), Steven Pinker (2002), and Levitt and Dubner (2006, p.157). In a 2018 interview with BBC World News HARDtalk, Plomin (2018) claimed that “(W)ith the exception of being in an abusive environment …, the people (including schools and jobs) who raised an individual would make little difference.”

Diametrically opposite to Arthur Jensen was the view of Claude Fischer and colleagues (e.g., 1996). They regarded a cognitive test such as the Armed Forces Qualification Test (AFQT) as a test of academic performance from school learning, which was heavily influenced by SES context. Positioned between the two extremes are those who considered cognitive ability a composite outcome that was shaped by both natural endowment and environmental conditions (e.g., Jencks et al. 1979; Scullin et al. 2000; Winship and Korenman 1997).

Regardless of their position on the role of heritance for cognitive ability, a fundamental weakness in all previous work is the lack of DNA data or data measured at the molecular level. As a substitute, biometric studies based on genetic relatedness of twins and sibling were used to estimate the heritability of cognitive ability. Heritability is the proportion of the total variance in a trait due to genetic factors. Under an array of assumptions that are difficult or impossible to satisfy (e.g., Nisbett 2009, Chapter 2), biometric analysis estimates the level of importance of unidentified genetic factors relative to that of non-genetic factors in a trait. It cannot estimate the effects of genotype side by side with the effects of SES context. Biometric studies yielded a heritability estimate that ranges between 0.2 to 0.8 (e.g., Bouchard 1998; Velden 1997) and a shared non-genetic influence that quickly reduces to zero after childhood (Bouchard 1998; Jensen 1997).

The policy arguments such as those made by Jensen (1969) and Herrnstein and Murray (1994) were based on a loose extrapolation of these biometric estimates. Persuasive evidence for policy arguments regarding individuals must be based on necessary genotype and social-contextual data at individual level. Such analysis cannot be carried out when only biometric data are available. Without such empirical work linking genotype to cognitive ability at individual level, heritability estimates can be easily loosely and incorrectly interpreted.

With the advent of DNA data and genome-wide association studies (GWAS) on outcomes closely related to cognitive ability, analysis becomes feasible that explicitly incorporates genomic measures. In this article, we report findings from a study that investigates how human genome and environment, especially SES context, jointly and interactively shape verbal intelligence.

## BACKGROUND

### Cognitive Ability and Life Outcomes

Cognitive ability has a strong positive association with academic performance in school. Examining six longitudinal studies, Jencks et al. (1979, p.102) concluded that the correlational association between the two ranged from 0.40 to 0.63. Using a large, nationally representative and prospective data source of more than 70,000 British students, Deary et al. (2007) showed that cognitive ability measured at 11 years old was a strong predictor of school achievement measured by national examinations on 25 subjects at 16 years old.

The positive association between cognitive ability and educational attainment is also evident. In a study that assembled eight large datasets including several massive state-wide datasets in the United States, Taubman and Wales (1972, p 20) reported a strong positive association between cognitive ability and continuing to college. While about 40-50% of the high school graduates at the 50th percentile of an ability test continued to college, the percentage continuing to college at the 90th percentile reached 80-90% in late 1950s and 1960s when the U.S. higher education expanded. Data from the Wisconsin Longitudinal Study (WLS) showed a positive association between an ability test score and length of schooling (Hauser 2010, p103).

The effect of cognitive ability on later life outcomes becomes complicated when the role of education is taken into consideration. Drawing data from the National Longitudinal Study of Youth (NLSY), Scullin et al. (2000) investigated the predictive power of AFQT administered when the respondents were 15 to 17 years old, together with educational attainment and race. They reported that although AFQT was predictive of labor market outcomes such as personal incomes, hourly wage, and occupational socioeconomic index (SEI), the predictive powers were substantially reduced when years of schooling was added to the models. The analysis by ethnicity showed that cognitive ability had a much smaller effect and educational attainment had a much larger effect for African Americans than for white Americans. Examining about 10,000 Wisconsin high school students followed since their graduation in 1957, Hauser (2010) concluded that the associations of cognitive ability with job performance, occupational prestige, income, and wealth largely disappeared once levels of schooling were adjusted.

The relationship between ability and personal health resembles that between ability and labor market outcomes. When family SES is adjusted and self educational attainment is not adjusted, ability proved to be a strong predictor of general health conditions. A 50-year Scottish longitudinal study on more than 27,000 adults reported that low cognitive ability at 11 years old was strongly associated with higher mortality before age 65 after adjusting for social class and deprivation category (Gruer, Hart and Watt 2017). Using the NLSY-79, Wraw et al. (2015) showed that AFQT was predictive of an array of health indicators measured at age 50 including three overall measures of general health, nine measures of diagnosed illness conditions, and two measures of self-reported conditions after family SES in childhood was adjusted. Once the respondents’ own SES in adulthood including education, income and occupation were adjusted, ability remained a significant predictor for the three overall measures of general health, but for many specific illness conditions, the effects sizes were reduced markedly.

### SES and Other Environmental Roots of Cognitive Ability

A body of evidence exists for the importance of SES context for cognitive ability. Among social-contextual factors, formal schooling is commonly viewed as the most direct and essential determinant of cognitive ability. Tests of cognitive ability are hardly meaningful out of the context of modern education. Dutch children’s schooling was delayed by the Nazi regime during WWII and these children’s IQ were seven points lower on average than those who went to school at regular ages after the war (DeGroot 1948; Nisbett 2009, p.41). In a longitudinal study of the effects of family SES, racial mix, and summer breaks on children’s mathematics achievement, Entwisle and Alexander (1992) concluded that “when school is in session, poor children and better-off children perform at almost the same level. Schools seem to be doing a better job than they have been given credit for (pp.82.)” They demonstrated the effect of schooling by showing the loss in mathematics scores after every summer break on the part of children in poverty relative to children not in poverty.

Through a natural experiment, Cahan and Cohen (1989) reported a schooling effect distinct from the effect of biological age on an IQ test score that included measurement of fluid intelligence. The study compared fourth graders and fifth graders who were essentially of the same age. Winship and Korenman (1997) reviewed the literature that estimated the effect of education on cognitive ability and carefully reanalyzed the NLSY data used by Herrnstein and Murray (1994). They concluded that each year of education increased 2.7 points of IQ units. Estimating the effect of education on cognitive ability is challenging because the two are intertwined. While academic training in school contributes to cognitive ability, cognitive ability is also related to how fast and efficiently academic subjects are absorbed and to eventual educational attainment. In this project, we estimated the effect of education on verbal ability measured at Wave 3 in Add Health after controlling for the exact same version of verbal ability administered at Wave 1. A similar strategy was used when estimating the effect of education on cognitive ability in the absence of DNA data (Winship and Korenman 1997, p 220).

Socioeconomic status is a composite concept with multiple dimensions that comprises family financial resources, knowledge, social connections, and the larger social context. SES is typically measured by parental income, education and occupation, one vs two biological parents in the household, sibship size, quality of neighborhood, and quality of school (e.g., Braveman et al. 2005; Link and Phelan 1996). In a meta-analysis of adoption studies of IQ, Locurto (1990) showed that adoption into high SES families raised the adopted children’s IQ by 10-12 points. These adoptive children tended to come from low SES families. Entwisle and Alexander (1992) concluded that family SES status had a larger impact than the interruption of schooling from summer breaks on cognitive test scores in the context of elementary education.

Mechanism studies explained why SES context makes a difference in cognitive test scores. Guo and Harris (2000) investigated the mechanisms through which family SES affected children’s cognitive ability. Using data from the NLSY, they showed that the effect of family SES was completely mediated by the intervening mechanisms measured by the latent factors of cognitive stimulation in home, home physical environment, parenting style, and child health at birth. Cognitive ability in that study was measured by four Peabody tests including a reading recognition test, a reading comprehension test, and a mathematics assessment test, and PPVT. Hart and Risley (1995) observed children in 42 families for one hour per week for two and a half years and their calculation suggested that large differences existed in the total number of words heard by children from birth to age four across professional families (45 millions), working class families (25 millions), and families in poverty (13 millions). Using the children of the NLSY79, Farkas and Beron (2004) showed that by 36-month of age, large gaps in vocabulary already emerged across social classes and racial/ethnic groups, and the gaps were not closed afterwards.

Macro historical trend of intelligence tests provides another source of evidence for environmental influences on cognitive ability. The well-known Flynn effect documented the historical rise in intelligence scores (e.g., Flynn 1987). The rise was interpreted as a result from a societal change that placed a greater value on rational reasoning in more recent time (Flynn 2007) as well as general improvements in education, nutrition and health (Flynn 2009).

Researchers have long examined the connections between problem behaviors and academic potential (e.g., Hinshaw 1992; Kessler et al. 1995; Malinauskiene et al. 2011; McLeod and Kaiser 2004). Our full models also tested the effects of binge drinking, marijuana use, smoking, and serious delinquency.

### Genomic Roots of Cognitive Ability

For decades, the genomic influences on cognitive ability were estimated via biometric studies. The estimated heritability has a vast range, depending on factors including sample size, the environment associated with the particular population from which the analysis sample is drawn, non-additive genetic variance, the assumptions of assortative mating and equal environment, and whether the data are based on a single genetic-relatedness such as adoptive-apart data or more than one genetic-relatedness such as the classic twin data (e.g., Bouchard and McGue 1981; Plomin and Loehlin 1989; Velden 1997). Heritability estimates, however, cannot be used to investigate the effects of environmental and genomic roots at the individual level.

The efforts linking DNA variation to cognitive ability began in earnest in the early 2000. By then, it was evident that intelligence was a complex trait subject to the influences of a large number of genes each with a tiny effect (Glazier, Nadeau and Aitman 2002). The challenge to find specific genes for cognitive ability was enormous. A human genome consists of millions of genetic variants. Testing whether each one of them predicts ability and setting the critical value for significance at the level of 0.05 would by chance generate a huge number of false positive findings. Although the genome is large, it is finite. The solution was to set a stringent critical value of 5×10^−8^ for significance and to request a replication of a discovered genetic variant in an independent data source. Initial successes of these genome-wide association studies (GWAS) employing several thousands of individuals were performed on human traits such as type 2 diabetes and body mass index (BMI) (Frayling et al. 2007; Zeggini et al. 2007). The number of GWAS-identified genetic loci were small, but tended to be replicated.

It soon became clear that by far the single most important factor in GWAS was sample size. The most readily available large samples were for anthropometric measures such as human height and body mass index (BMI). The 2010 (Allen et al. 2010) and 2014 (Wood et al. 2014) height GWAS had a sample size of 183,727 and 253,288 and identified 180 and 423 genetic loci, respectively. A 2017 study based on 711,428 individuals discovered additional 83 rare (minor allele frequency [MAF]<1%) and low-frequency (1%<MAF<5%) coding variants associated with height (Marouli et al. 2017). The 2010 (Speliotes et al.) and 2015 (Locke et al.) GWAS on BMI, respectively, identified 32 and 97 genetic loci. The later round of GWAS generally included all the observations used in a previous round of GWAS. For example, the 2015 BMI GWAS included all the individuals used in the 2010 BMI GWAS. It was of no surprise that a later GWAS tended to replicate the loci identified in an earlier GWAS.

The prospect of discovering specific genes for intelligence appeared more daunting than the genes for anthropometric measures. Human intelligence seems to be much more complicated than height and weight. Even how intelligence is measured can be highly contentious. The candidate-gene approach was called into question when Chabris et al (2012) reported that they could not replicate the previously published associations between 12 specific genetic variants and cognitive ability. An early GWAS based on 3,511 individuals targeting at intelligence failed to find any signal that was genome-wide significant at *P*<10×5^−8^ (Davies et al. 2011).

A 2013 GWAS study of education based on a discovery sample of 101,069 obtained three SNPs with genome-wide significance, one of the three associated with years of education and the other two with a college degree (Rietveld et al. 2013). The effect sizes of these SNPs were about one tenth of those of height and weight. A polygenic score (PGS) constructed from all common SNPs explained about 2% of the variance in both educational attainment and cognitive function. Two subsequent GWAS of educational attainment in 2016 (Okbay et al.) and 2018 (Lee et al.) assembled 293,723 and 1.1 million individuals, and reported 74 and 1,271 independent genome-wide significant SNPs associated with years of education, respectively. The PGS constructed from all common SNPs in the latest GWAS (Lee et al. 2018) obtained an *R^2^* of about 12% using data from Add Health. Many of the genetic loci were implicated in biological pathways that played a role during prenatal brain development. The estimated genetic overlap or the shared genetic influences between years of education and cognitive performance was about 70% (Supplementary Table 3.1, Okbay et al. 2016), suggesting that the education-based GWAS-identified genetic variants ought to be reasonable predictors of cognitive ability.

Two successive GWAS of cognitive ability in 2017 (Sniekers et al.) and 2018 (Savage et al.) employed 78,308 and 269,867 individuals, respectively. The 2018 GWAS identified 205 independent genome-wide significant loci, which included the 15 variants identified in the 2017 GWAS. The PGS based on the 2018 GWAS obtained an incremental *R*^2^ of about 5% for cognitive ability. The study used a variety of tests measuring cognitive ability because the analysis sample was assembled from more than a dozen cohorts. These tests either measured fluid intelligence or were used to calculate the Spearman’s g. The genetic correlation between intelligence and education was estimated again to be about 0.70 with *P*=2.5×10^−287^ by the whole-genome LD score regression (Bulik-Sullivan et al. 2015), which computed the correlation between the genetic influences behind intelligence and those behind education. The GWAS-identified genes were mostly expressed in brain tissues. In the present analysis, the latest findings of GWAS for intelligence and the findings of GWAS for educational attainment were used to construct the genomic measures to be included in models predicting verbal ability.

In addition to the two genomic measures of educational attainment and intelligence, we tested a number of other genomic measures. Previous work documented the association of academic achievement and educational attainment with obesity (Della Bella and Lucchini 2015; Roskam et al. 2010), general health (Ickovics et al. 2014), health behavior (McLeod and Kaiser 2004), personality (Lynn and Gordon 1961; McKenzie, Taghavi-Khonsary and Tindell 2000; Phillips and Endler 1982), stress (McEwen 2000), and brain size (Pietschnig et al. 2015). To assess the effects of these potential predictors of cognitive ability, we included a number of PGSs that underlie these phenotypic predictors. These PGSs consisted of those based on GWAS for general health-related birthweight (Okbay et al. 2016), BMI (Locke et al. 2015), number of cigarettes smoked per day (Furberg et al. 2010), head circumference (Taal et al. 2012), and each of the big five personality traits (agreeableness, conscientiousness, extraversion, neuroticism, and openness) (de Moor et al. 2012). We included the PGSs that underlined these traits rather than observed traits themselves to reduce the difficulty of interpretation. For example, an observed BMI at 15 years old is likely a result of a host of genetic and environmental influences the individuals experienced up to that point in life. Some of these environmental influences are likely to be correlated with those that affect cognitive development.

### Gene-Environment Interactive Effects

Gene-by-environment (GE) interaction refers to the interdependence between an environmental effect and a genotype effect. In the present setting, our GE-interaction hypothesis predicts that a favorable SES context enhances the effects of ability PGSs. The reasoning goes that a favorable SES context would help realize or even raise the cognitive ability of an individual whereas individuals in an unfavorable SES context would have a verbal ability score below that predicted by his or her ability PGSs. GE-interaction analysis is one of the most valuable tools of social scientists who strive to understand the interplay between genetics and SES context. Previous GE-interaction work on cognitive development using biometric data generated evidence for our hypothesis (Guo and Stearns 2002; Turkheimer et al. 2003). The present project tested the hypothesis by taking advantage of rich SES-contextual measures and DNA-based PGSs in Add Health.

### Genetic Analysis Using Race/Ethnicity Samples

To include or exclude ethnic minorities in our analysis is an important and difficult issue (Hindorff et al. 2018). On one hand, the diversity requirement of the National Institutes of Health (NIH) has been a top priority since the 1993 Revitalization Act that requires that all federally funded clinical research prioritize the inclusion of women and minorities. Compared with some other large longitudinal studies in social sciences such as the Health and Retirement Survey and the Wisconsin Longitudinal Study, Add Health is particularly well-suited for a study that includes ethnic minorities because of Add Health’ deliberate inclusion of a large number of African and Hispanic Americans from the get-go.

On the other hand, it is well-known that the GWAS findings do not apply to ethnic minorities as well as they do to European Americans. Notable genomic differences across individuals from different continents have been known since 1990s (Cavalli-Sforza, Menozzi and Piazza 1996). The recent 1000-genome project shows that although more than 70% of genomic variation are common across continental populations, individuals of African descent have about 20% more unique genetic variants than individuals of European ancestry (Consortium 2015).

The published GWAS studies were mostly based on individuals of European descent including the GWAS that identified the genetic loci associated with educational attainment and cognitive ability. In the over 3,100 published GWAS by 2017, about 19% of individuals included in GWAS were non-European and the large majority of the 19% were East Asians (MacArthur et al. 2017). The partial overlap between the European genetic variants and those of Africans and Hispanics implies that the current GWAS must have missed a substantial number of genetic loci among the ethnic minorities that are associated with educational attainment and intelligence. This, in turn, suggests that although the PGSs based on current GWAS can still be significantly predictive of cognitive ability among ethnic minorities, their predictive power is expected to be lower in the minority samples than in the European samples (Duncan et al. 2018).

To take advantage of Add Health’ diversified data source, we kept the samples of ethnic minorities in the analysis. To characterize the potential differential predictability of the PGSs, we conducted the analysis by ethnicity. To minimize the noises from small samples, we combined the ethnic groups in analysis when the predictability of the PGSs were similar across the ethnic groups. Preserving statistical power is crucial because other than genomic differences small sample sizes could also lead to an idiosyncratic set of findings.

### Verbal Ability

At Waves I and III, Add Health implemented an abridged version (AHPVT or PVT) of the Peabody Picture Vocabulary Test-Revised (PPVT-R). Our analysis used the percentile rank of PVT ranging 0-100 that was translated from the raw PVT score. PPVT was first published in 1959 and has been revised three times (Dunn and Dunn 2007). Psychological literature considers PPVT an estimate of verbal intelligence (Beres, Kaufman and Perlman 1999; Campbell and Dommestrup 2010). The correlations between PPVT and full-scale intelligence tests were found to be between 0.40 to 0.60 (Beres, Kaufman and Perlman 1999). Carvajal et al. (1993) empirically estimated the correlation between the Wechsler Intelligence Scale for Children-III and the Peabody Picture Vocabulary Test-Revised among children enrolled in Grades 3, 4, and 5. They obtained statistically significant correlations of .75, .76, and .60, between PPVT and the Wechsler Vocabulary subtest scaled scores, the Wechsler Verbal scores, and the Wechsler Full Scale IQ scores, respectively. Bell et al. (2001) compared PPVT-III and Wechsler Adult Intelligence Scale-Third Edition (WAIS-III) using a sample of 40 adults aged 18-41 with a mean of 22 years. Their results showed that PPVT-III was statistically significantly correlated with the WAIS-III full-scale IQ (FSIQ, 0.40) and verbal IQ (VIQ. 0.46), and that PPVT-III is uncorrelated with performance IQ (VIQ). Compared with full-scale intelligence tests, PPVT is easier-to-use and time-saving. These are decisive advantages in a setting of a large survey like Add Health that includes a large number of survey items besides a verbal ability test.

## DATA, MEASURES AND METHODS

### Data Source

We used data from the National Longitudinal Study of Adolescent to Adult Health (Add Health) (http://www.cpc.unc.edu/projects/addhealth/), which is an ongoing longitudinal study of a nationally representative sample of more than 20,000 adolescents in grades 7-12 in 1994-95 in the United States who have been followed for more than 20 years. Over the years, Add Health has conducted one in-school survey in 1994-1995, and five in-home interviews in 1994-1995 (Wave 1), 1996 (Wave 2), 2001-02 (Wave 3), 2008 (Wave 4), and 2016-8 (Wave 5). Add Health includes more than 3,000 individuals who are identical twins, fraternal twins, full siblings, and half siblings. Add Health has a multiracial and multiethnic sample. The original purpose of Add Health was to understand the causes of health, health behavior, and educational development with special emphasis on the role of social context at the levels of family, neighborhoods, and communities.

We started with a sample of 9,975 individuals for whom GWAS measures were available. Our final analysis sample consisted of 8,078 individuals. Excluded were those without a measure of verbal ability (1,859) and those (38) who self-identified as Native Americans. A total of seven individuals had missing values in one of the following covariates: whether the respondent was in school before the interview, binge drinking, and marijuana use and smoking. These seven cases were replaced by the respective sample means. Fifty-one individuals with missing values in race/ethnicity were reclassified into the category of Hispanics. Missing values in SES variables were coded into a separate category.

In January 2015, Add Health completed genome-wide genotyping on the Wave IV participants who consented to archive their DNA for future studies. Of the 15,701 respondents interviewed, 12,200 of the eligible respondents agreed to archive their DNA for future analysis “related to long term health.” The consent was largely uniform across racial/ethnic groups and yielded more than 12,000 samples for genome-wide genotyping. Add Health utilized two Illumina platforms for GWAS: the Illumina Human Omni1-Quad BeadChip at first and the Illumina Human Omni-2.5 Quad BeadChip at a later time. The two platforms utilized tag SNP technology to identify and include, respectively, >1.1 million and 2.5 million genetic markers, which were derived from phases 1-3 of the International HapMap Project (Altshuler et al. 2005) and the 1,000 Genomes Project (1KGP) (Altshuler et al. 2010).

### Measures of Verbal Ability, SES Context and Other Covariates

*Verbal Ability* was measured by an abridged version (PVT) of the Peabody Picture Vocabulary Test-Revised (PPVT-R) implemented twice at Waves 1 and 3 at Add Health. PVT included 87 or half of the items of PPVT-R. Our analysis used the PVT percentile rank as the outcome variable. The variable was constructed by computing the unsmoothed percentile rank for the same-age peers at each Add Health Wave. The resulting percentile rank was comparable across age groups at the same Wave.

#### SES Context

Mother’s and father’s education were, respectively, coded into a four-category categorical variable with less than high school, high school graduation and some college, at least college degree, and missing. Mother’s and father’s occupation originally had 16 categories. They were combined into five categories of none and other, which were two of the original categories of Add Health; manual or blue collar; sales, service, or administrative; professional or managerial; and missing. Household income was total family income obtained from the parental questionnaire at Wave 1 in 1994. Household income was coded into a six-category variable with cutoff points at 20^th^, 40^th^, 60^th^, and 80^th^ percentile plus a category of missing. Family structure was measured by a dichotomous variable taking the value of one if the respondent lived with two biological parents and zero if the respondent was from a household of a single parent, a stepparent, adopted families, and foster homes at Wave 1. Sibship size was the number of siblings living in the household at Wave 1.

To capture neighborhood disadvantage, we followed the approach used by Wodtke, Harding and Elwert (2011). Neighborhood disadvantage was measured by the first principal component from a principal component analysis of six neighborhood measures that included the proportion of households living below the poverty line, the proportion of adults who were unemployed, the proportion of female-headed households, the proportion of adult residents without a high school diploma, the proportion of residents with a college degree, and the proportion of workers holding managerial or professional jobs. In school was coded as one if the respondent was in a school session when interviewed or if the respondent was in school in the past school year when interviewed in a summer break; and in school was coded as zero otherwise.

In generating covariates, we took advantage of the longitudinal design of Add Health. Whenever possible, we created time-varying covariates using repeated measures of Add Health. For example, we generated two measures of neighborhood disadvantage derived from two rounds of principal component analysis using data at Waves 1 and 3, respectively. These two measures of neighborhood disadvantage were included longitudinally in our analysis. Similarly, parental education and occupation were also measured at Waves 1 and 3 and included as time-varying covariates in analysis.

#### General Health and Health Behaviors

Self-reported health measured general health of the respondents at Waves 1 and 3 and was based on an answer to the question of "(I)n general, how is your health?” The answer had five categories of 1=excellent, 2=very good, 3=Good, 4=fair, and 5=Poor. To facilitate interpretation, the variable was reversely coded so that 1=Poor, 2=Fair, 3=Good, 4=Very good, and 5=Excellent, and treated as a continuous variable in analysis. Binge drinking was measured by the number of days the respondents drank 5 or more drinks in a row throughout a year at Waves 1 and 3. The survey response categories of binge-drinking every day or 3 to 5 days a week, 1 or 2 days a week, 2 or 3 days a month, 3 to 12 times in the past 12 months, 1 or 2 days in the past 12 months, and never were coded into 6-categorical category variable with never as the reference category. Marijuana use was measured by the number of times the respondents used marijuana during the past 30 days at Waves 1 and 3. Those who skipped the question legitimately were coded as 0 and the variable had a range of 0 to 900. Then the variable was recoded into categories of 0, 1-5, 6-15, 16-30, and 31 or more. Smoking was measured by the number of cigarettes smoked per day over the past 30 days. Those who skipped the question legitimately were coded as 0 and the variable ranged from 0 to 100. The variable was recoded into categories of 0, 1-5, 16-30, and 31-100.

Serious delinquency was constructed from a delinquency scale using 12 questions at Waves 1 and 3. The scale was similar to those widely used in delinquency and criminal behavior research (Thornberry and Krohn 2000). The 12 questions were about physical fighting that resulted in injuries needing medical treatment, use of weapons to get something from someone, involvement in physical fighting between groups, shooting or stabbing someone, deliberately damaging property, pulling a knife or gun on someone, stealing amounts larger or smaller than $50, breaking and entering, and selling drugs. Respondents were asked to report how many times they had been engaged in the delinquent behaviors in the past 12 months. The answers are none, 1 or 2 times, 3 or 4 times, and 5 or more times. We recoded them into 0, 1, 2, and 3, respectively, for each item. The scores were then summed up and divided it by twelve. The results were rounded up and coded into categories of none, 1 or 2 times, 3 or more times, and missing. *Demographic variables* included gender, age at Waves 1 and 3, and race and ethnicity. US born and speaking English at home were used to capture the immigration status of the respondent.

To address the common difficulty of lack of power in gene-environment interaction analysis, we constructed an SES Composite (Yang et al. 2017). The SES composite included parental education, parental occupation, household income, sibship size, neighborhood disadvantage, and in school. Parental education was the higher value of mother’s and father’s education. Parental occupation was the higher occupational prestige score of the two parents. The categories of household income and sibship size were treated as continuous. The coding of neighborhood disadvantage and in school was unchanged. The recoded variables were individually standardized first. The SES composite was the mean of the six standardized items. The genomic composite score was constructed similarly from the two ability PGSs.

### Genomic Measures

The genomic measures of educational attainment and cognitive ability were based on the 2018 GWAS of years of schooling (Lee et al. 2018) and the 2018 GWAS of intelligence (Savage et al. 2018), respectively. In the case of educational attainment, the GWAS separately regressed each of a large number of single nucleotide polymorphisms (SNPs) on years of schooling where SNPs were a particular type of genetic variables taking values 0, 1, or 2. The value represented the number of risk alleles at the genetic loci for the phenotype. The GWAS obtained one *β* from each SNP. The study identified 1,271 *β*s significant at the genome-wide level of 5×10^−8^. To tap all the predictability of a GWAS, a polygenic score (PGS) for individual *i* is often constructed using all *β*s from a GWAS as weights for the observed risk alleles (Χ_*ij*_) for this individual *i*: 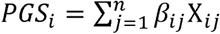, where *i* indexes individual, *j* indexes SNP, and *n* stands for the total number of SNPs used in the calculation. The total number of SNPs included in the calculation were a subset of the GWAS SNPs. The subset was obtained from pruning and clumping the original GWAS SNPs. The SNPs in the subset were uncorrelated or weakly correlated with one another. Pruning is used to remove some of the highly correlated SNPs and clumping is used to keep only one SNP in a section in which the SNPs are highly correlated. When *n* is equal to the entire set of SNPs, it is equivalent to setting *P*=1, where *P* is the critical value in a SNP regression.

To facilitate interpretation, a PGS is typically standardized into a Z score: 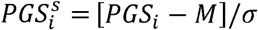, where M is the mean PGS averaged over all individuals in the sample and σ is the standard deviation of the PGS. When 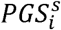 is included in a regression model predicting verbal ability, its coefficient can be interpreted as the effect of one standard deviation of the PGS. Thus, the standardized PGSs are a way to estimate and interpret the overall genomic influence on a phenotype. We have similarly constructed standardized PGSs for intelligence, BMI, head circumference, cigarette smoke per day, birthweight, and the Big Five personality traits of agreeableness, conscientiousness, extraversion, neuroticism, and openness. In the rest of this article, we used *PGS_i_* to represent 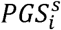 for simplicity.

## ANALYTICAL STRATEGIES

Our Add Health data had a hierarchical structure that induces correlation in the outcome variable and that must be addressed statistically in a regression setting (e.g., Diggle, Liang and Zeger 1994). One source of the hierarchy was due to the inclusion of the two measures of verbal ability per individual. The other source of the hierarchy originated from the study design of Add Health, which included a genetic-informative sample consisting of full siblings, dizygotic (DZ) twins, monozygotic (MZ) twins, and other related individuals. When these siblings were first collected, they were the source of information on inheritance for biometric studies. In the present study, they were complications that needed to be addressed. In our analysis, we addressed the hierarchy of the data using mixed regression models (Raudenbush and Bryk 2002; Searle 1971; Searle, Casella and McCulloch 1992), which are also referred to as random-effects models or multi-level models. The mixed models have long been established in the statistical literature for data that are not independent. We implemented two types of mixed models written below in the form of multilevel models. The first was a three-level model that addressed the repeated measures of verbal ability in addition to the sibling clusters:

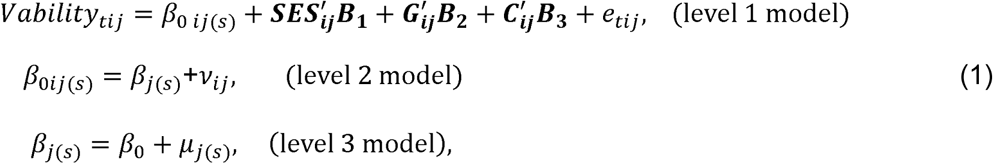

where *Vability* stands for verbal ability; the subscripts *t*, *i*, *j*, and *s* index Add Health Wave, individual, sibship and type of sibship, respectively; ***SES***, ***G*** and ***C*** are, respectively, vectors of SES measures, PGSs, and other variables including demographic indicators and principal components for addressing population admixture; ***B_1_***, ***B_2_***, and ***B_3_*** are vectors representing the effects of these observed variables; and *e_tij_, v_ij_*, and *µ_j(s)_* are random effects at the level of Add Health Wave, individual and sibship, respectively. We estimated models that distinguished different types of sibship and models that did not make that distinction, which was equivalent to ignoring (*s*). The two sets of estimated coefficients of observed variables were essentially identical. We only presented estimates from the models that did not make the distinction.

Our mixed model that conditioned on Wave-1 verbal ability was a two-level model that used verbal ability at Wave 3 as the dependent variable and estimated the effects of the same set of predictors while controlling for verbal ability at Wave 1:

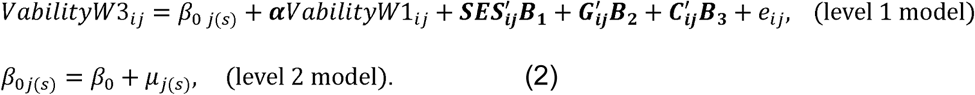

Both (1) and (2) were random-intercept models. Conditional on the random intercepts at the individual level and the level of sibling clusters, the siblings and repeated measures were assumed to be independent.

Population admixture or population stratification is a major concern in genetic association studies. Population groups separated over the past 50,000-100,000 years are likely to have developed private genetic variants that differ across population groups and that are unrelated to cognitive ability. If these population groups differ in ability test scores for social-contextual and other environmental reasons and if these private variants are not controlled properly, the association between the variants and test scores could be erroneously interpreted as causal. The error can be avoided by the common practice of including the ten or so largest principle components (PCs) in the regression that links genetic variants to a phenotype (Price et al. 2006). The PCs represent ancestral genetic differences among population groups and are highly correlated with self-reported race/ethnicity. GE interaction terms can be added to models (1) and (2) readily.

## FINDINGS

Table 1 showed the means and the standard deviations of the continuous variables, and the percentage distribution of categorical variables used in the analysis for whites, blacks and Hispanic blacks, Asians, Hispanic whites, and all ethnicities.

**Table 1.** Descriptive statistics: Add Health by Ethnicity

| Variable | White<br>(N=4,860) |  | Black & Hispanic<br>Black<br>(N=1,677) |  | Asian<br>(N=438) |  | Hispanic<br>White<br>(N=1,103) |  | All<br>(N=8,078) |  |
| --- | --- | --- | --- | --- | --- | --- | --- | --- | --- | --- |
|  | Mean or<br>% | S.D. | Mean or<br>% | S.D. | Mean or<br>% | S.D. | Mean or<br>% | S.D. | Mean or<br>% | S.D. |
| **PVT Percentile Score (Verbal Ability)** | 58.0 | 26.3 | 35.7 | 28.0 | 49.0 | 30.8 | 40.0 | 29.3 | 50.4 | 29.0 |
| **Ability Polygenic Scores (PGS)** |  |  |  |  |  |  |  |  |  |  |
| PGS for education | 0.4 | 0.5 | -1.5 | 0.9 | 0.6 | 0.6 | 0.2 | 0.6 | 0.0 | 1.0 |
| PGS for cognitive ability | 0.5 | 0.8 | -1.2 | 0.7 | -0.5 | 0.6 | 0.2 | 0.8 | 0.0 | 1.0 |
| **SES Context** |  |  |  |  |  |  |  |  |  |  |
| Mother's education |  |  |  |  |  |  |  |  |  |  |
| Less than high school | 11.0 | - | 13.0 | - | 12.3 | - | 36.5 | - | 15.0 | - |
| High school/some college | 59.0 | - | 53.8 | - | 36.8 | - | 43.2 | - | 54.6 | - |
| ≥college graduation | 27.3 | - | 27.7 | - | 43.2 | - | 12.4 | - | 26.2 | - |
| Missing | 2.7 | - | 5.5 | - | 7.7 | - | 8.0 | - | 4.3 | - |
| Father's education |  |  |  |  |  |  |  |  |  |  |
| Less than high school | 10.6 | - | 13.2 | - | 11.7 | - | 37.6 | - | 14.9 | - |
| High school/some college | 56.9 | - | 54.5 | - | 35.0 | - | 40.1 | - | 52.9 | - |
| ≥college graduation | 26.6 | - | 24.9 | - | 42.5 | - | 12.1 | - | 25.1 | - |
| Missing | 5.9 | - | 7.4 | - | 10.9 | - | 10.2 | - | 7.1 | - |
| Mother's occupation |  |  |  |  |  |  |  |  |  |  |
| None & other | 23.9 | - | 24.4 | - | 18.9 | - | 32.6 | - | 24.9 | - |
| Manual or blue collar | 15.0 | - | 21.2 | - | 17.2 | - | 26.1 | - | 17.9 | - |
| Sales/service/administrative | 26.3 | - | 18.9 | - | 24.4 | - | 20.0 | - | 23.8 | - |
| Professional or managerial | 29.9 | - | 31.5 | - | 34.1 | - | 16.6 | - | 28.6 | - |
| Missing | 4.9 | - | 3.9 | - | 5.3 | - | 4.8 | - | 4.7 | - |
| Father's occupation |  |  |  |  |  |  |  |  |  |  |
| None & other | 11.8 | - | 9.5 | - | 13.0 | - | 14.4 | - | 11.7 | - |
| Manual or blue collar | 36.3 | - | 25.5 | - | 33.4 | - | 43.2 | - | 34.9 | - |
| Sales/service/administrative | 6.4 | - | 3.0 | - | 10.4 | - | 3.3 | - | 5.5 | - |
| Professional or managerial | 26.0 | - | 12.0 | - | 30.6 | - | 13.4 | - | 21.6 | - |
| Missing | 19.4 | - | 50.1 | - | 12.6 | - | 25.7 | - | 26.3 | - |
| Household income |  |  |  |  |  |  |  |  |  |  |
| 0-20 percentile | 9.4 | - | 25.2 | - | 5.2 | - | 19.9 | - | 13.9 | - |
| 20-40 percentile | 12.2 | - | 15.4 | - | 8.9 | - | 18.0 | - | 13.5 | - |
| 40-60 percentile | 18.6 | - | 13.2 | - | 16.8 | - | 15.5 | - | 17.0 | - |
| 60-80 percentile | 21.4 | - | 10.9 | - | 15.1 | - | 12.5 | - | 17.7 | - |
| 80-100 percentile | 21.0 | - | 9.7 | - | 17.8 | - | 7.6 | - | 16.7 | - |
| Missing | 17.3 | - | 25.7 | - | 36.3? | - | 26.5 | - | 21.3 | - |
| Family Structure |  |  |  |  |  |  |  |  |  |  |
| 2 Biological Parents | 0.5 | - | 0.3 | - | 0.5 | - | 0.5 | - | 0.5 | - |
| Sibling size |  |  |  |  |  |  |  |  |  |  |
| No sibling | 4.1 | - | 3.8 | - | 4.6 | - | 3.9 | - | 4.1 | - |
| 1 to 2 sibling(s) | 55.6 | - | 33.7 | - | 54.9 | - | 41.1 | - | 49.1 | - |
| 3 to 5 siblings | 26.5 | - | 31.9 | - | 25.6 | - | 34.1 | - | 28.6 | - |
| 6 to 20 siblings | 13.6 | - | 30.4 | - | 14.9 | - | 20.7 | - | 18.1 | - |
| Other and Missing | 0.1 | - | 0.2 | - | - | - | 0.2 | - | 0.1 | - |
| Neighborhood Disadvantages | 0.0 | 1.0 | 0.0 | 1.0 | 0.1 | 1.2 | 0.0 | 0.9 | 0.0 | 1.0 |
| In School | 0.5 | 0.5 | 0.5 | 0.5 | 0.6 | 0.5 | 0.5 | 0.5 | 0.5 | 0.5 |
| **Demographics** |  |  |  |  |  |  |  |  |  |  |
| Age | 19.1 | 3.6 | 19.2 | 3.6 | 19.6 | 3.5 | 19.6 | 3.6 | 19.2 | 3.6 |
| Gender |  |  |  |  |  |  |  |  |  |  |
| Female | 0.5 | - | 0.6 | - | 0.5 | - | 0.5 | - | 0.5 | - |
| Male | 0.5 | - | 0.4 | - | 0.5 | - | 0.5 | - | 0.5 | - |
| Race & Ethnicity |  |  |  |  |  |  |  |  |  |  |
| White | - | - | - | - | - | - | - | - | 60.2 | - |
| Black | - | - | - | - | - | - | - | - | 20.7 | - |
| Asian | - | - | - | - | - | - | - | - | 5.4 | - |
| Hispanic | - | - | - | - | - | - | - | - | 13.7 | - |
| Immigration Status |  |  |  |  |  |  |  |  |  |  |
| US Born | 0.7 | - | 0.7 | - | 0.5 | - | 0.5 | - | 0.7 | - |
| Speaking English at Home | 1.0 | - | 1.0 | - | 0.7 | - | 0.6 | - | 0.9 | - |
| **General Health and Health Behaviors** |  |  |  |  |  |  |  |  |  |  |
| Self-reported General Health | 3.9 | 0.9 | 3.9 | 0.9 | 3.9 | 0.9 | 3.9 | 0.9 | 3.9 | 0.9 |
| Binge Drinking |  |  |  |  |  |  |  |  |  |  |
| 0 times a year | 55.4 | - | 79.7 | - | 67.2 | - | 61.0 | - | 61.9 | - |
| 1 to 12 times a year | 24.8 | - | 11.6 | - | 21.1 | - | 23.2 | - | 21.6 | - |
| 2 or 3 days a month | 8.4 | - | 3.4 | - | 4.7 | - | 6.4 | - | 6.9 | - |
| 1 or 2 days a week | 7.7 | - | 2.7 | - | 5.0 | - | 6.5 | - | 6.3 | - |
| 3 to 5 days a week/(Almost) Every day | 3.7 | - | 2.6 | - | 2.0 | - | 3.0 | - | 3.3 | - |
| Marijuana Use |  |  |  |  |  |  |  |  |  |  |
| 0 time in past 30 days | 79.7 | - | 82.6 | - | 86.0 | - | 81.7 | - | 80.9 | - |
| 1 to 5 times | 11.5 | - | 9.8 | - | 9.0 | - | 10.6 | - | 10.9 | - |
| 6 to 15 times | 3.1 | - | 2.9 | - | 1.8 | - | 2.5 | - | 2.9 | - |
| 16 to 30 times | 4.7 | - | 3.8 | - | 2.4 | - | 3.9 | - | 4.3 | - |
| 31 times and above | 1.1 | - | 1.0 | - | 0.9 | - | 1.4 | - | 1.1 | - |
| Smoking |  |  |  |  |  |  |  |  |  |  |
| 0 cigarettes per day | 63.3 | - | 81.7 | - | 75.2 | - | 76.8 | - | 69.6 | - |
| 1 to 5 cigarettes | 15.5 | - | 13.0 | - | 14.6 | - | 14.5 | - | 14.8 | - |
| 6 to 15 cigarettes | 12.8 | - | 3.8 | - | 7.8 | - | 6.1 | - | 9.7 | - |
| 16 to 30 cigarettes | 7.9 | - | 1.4 | - | 2.2 | - | 2.4 | - | 5.5 | - |
| 31 to 100 cigarettes | 0.6 | - | 0.2 | - | 0.1 | - | 0.3 | - | 0.5 | - |
| Serious Delinquency |  |  |  |  |  |  |  |  |  |  |
| None | 63.0 | - | 57.6 | - | 60.0 | - | 60.0 | - | 61.3 | - |
| 1 or 2 times | 34.6 | - | 37.1 | - | 36.3 | - | 35.4 | - | 35.3 | - |
| 3 or more times | 1.0 | - | 1.5 | - | 1.4 | - | 1.8 | - | 1.3 | - |
| Missing | 1.4 | - | 3.8 | - | 2.4 | - | 2.8 | - | 2.2 | - |
| **Other Polygenic Scores** |  |  |  |  |  |  |  |  |  |  |
| PGS for Birthweight | 0.6 | 0.5 | -1.5 | 0.7 | -0.5 | 0.5 | 0.0 | 0.6 | 0.0 | 1.0 |
| PGS for BMI | -0.7 | 0.5 | 1.4 | 0.5 | 1.1 | 0.5 | 0.3 | 0.7 | 0.0 | 1.0 |
| PGS for Head Circumference | 0.6 | 0.4 | -1.6 | 0.4 | -0.6 | 0.3 | 0.0 | 0.4 | 0.0 | 1.0 |
| PGS for Number of Cigarette Per Day | 0.5 | 0.7 | -1.1 | 0.8 | -0.9 | 0.7 | -0.2 | 0.8 | 0.0 | 1.0 |
| PGS for Agreeableness | 0.5 | 0.8 | -0.9 | 0.6 | -1.6 | 0.8 | -0.1 | 0.8 | 0.0 | 1.0 |
| PGS for Conscientiousness | 0.4 | 0.8 | -0.4 | 0.7 | -1.8 | 1.0 | -0.4 | 0.9 | 0.0 | 1.0 |
| PGS for Extraversion | -0.1 | 1.0 | 0.7 | 0.8 | -1.1 | 0.9 | -0.2 | 0.9 | 0.0 | 1.0 |
| PGS for Neuroticism | -0.3 | 0.9 | 1.0 | 0.7 | -0.3 | 0.8 | -0.1 | 0.9 | 0.0 | 1.0 |
| PGS for Openness | -0.6 | 0.7 | 0.9 | 0.6 | 2.0 | 0.8 | 0.3 | 0.8 | 0.0 | 1.0 |

Table 2 presented the coefficients and their standard errors of SES context from random-effects models of twice-measured verbal ability for whites, blacks and Hispanic blacks, Asians, Hispanic whites, the combined sample of whites, Asians, and Hispanic whites (the combined sample), and all ethnicities. The random-effects model was a 3-level random-effects model of verbal ability (Equation 1). Little surprise that the two random effects at the individual and family levels were highly statistically significant. The OLS R^2^ was estimated to range from 13.4% to 23.5% across the different samples.

**Table 2.**
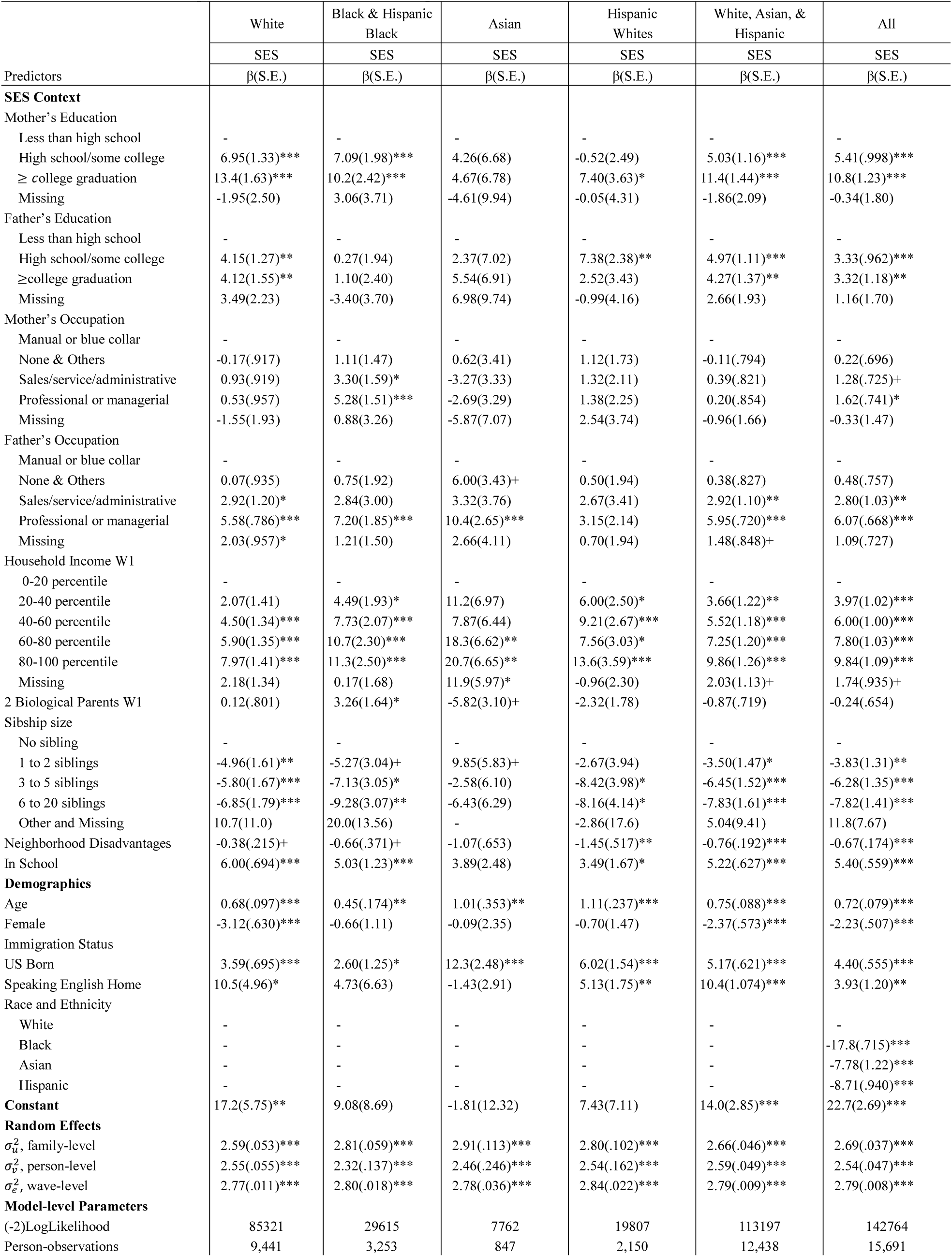
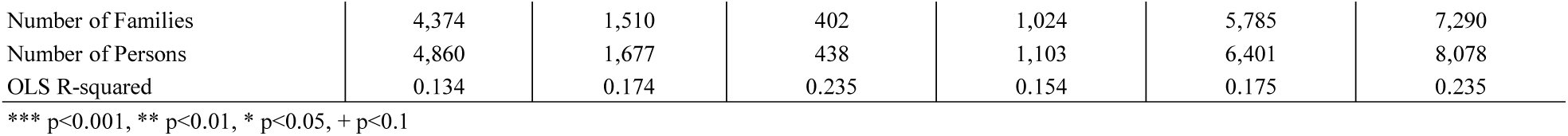
Random-effects models of twice-measured verbal ability: Coefficients (standard errors) of SES context in ethnicities-specific and -combined samples of Add Health

The models presented in Table 2 were traditional social-science model that did not consider genomic influences. SES context was important in all models though some differences were present across ethnic groups. Among whites, controlling for demographic factors, all social-contextual characteristics except for mother’s occupation significantly and simultaneously predicted verbal ability in directions consistent with expectation. For example, individuals whose mothers with at least a college degree scored 13.5 points higher on the verbal ability test than those whose mothers with an education less than high school. Those whose fathers with at least a college degree scored 4.1 points higher than those whose fathers with an education less than high school. Those whose fathers holding a professional and managerial job was about 5.58 points higher than those whose fathers holding a manual and blue-collar job. Individuals living in a household in the top 20% income group scored about eight points higher than those in the lowest 20% income group. Individuals living in a household with 3 to 5 siblings scored 5.8 points lower than those who were the only child in a family. An increase in one standard deviation in the index of the neighborhood disadvantage was associated with a decrease of 0.4 point of verbal ability score at a level of 10 percent. Those who were in a school session when taking the verbal ability test scored 6.0 points higher than those who were not in a school session. The coefficients estimated among Asians and Hispanic whites tended to be broadly similar to those for whites. This observation was the rationale for combining whites, Asians, and Hispanic whites. In the black sample, the effects of father’s education were much weaker than those in the white sample and the effects of mother’s occupation were much stronger than those in the white sample. The estimates in the Asian sample tended to be less statistically significant than those in the white sample.

Table 3 presented the coefficients and their standard errors of ability-related PGSs from random-effects models of twice-measured verbal ability for the same set of samples as in Table 2. The PGSs were standardized in the overall sample so that the coefficients across ethnicities can be compared. Among white respondents, one standard deviation of the education PGS was associated with 9.56 points of verbal ability or 9.56/50=19% of the mean of verbal ability. The education PGS was significantly predictive of verbal ability in all estimated models and its coefficients were similar in size except for the black sample in which the coefficient was much smaller. The IQ PGS significantly predicted verbal ability in all samples except the black sample. The ten principal components were mostly significant in the overall sample and the combined sample because these PCs were highly correlated with race and ethnicity. These results established the importance of the two PGSs.

**Table 3.** Random-effects models of twice-measured verbal ability: Coefficients (standard errors) of ability-related PGSs in ethnicities-specific and -combined samples of Add Health

|  | White | Black & Hispanic<br>Black | Asian | Hispanic | White, Asian, &<br>Hispanic | All |
| --- | --- | --- | --- | --- | --- | --- |
|  | PGS | PGS | PGS | PGS | PGS | PGS |
| Predictors | $\beta$(S.E.) | $\beta$(S.E.) | $\beta$(S.E.) | $\beta$(S.E.) | $\beta$(S.E.) | $\beta$(S.E.) |
| **Ability PGSs** |  |  |  |  |  |  |
| PGS for Education | 9.56(.644)*** | 1.74(.730)* | 7.26(2.33)** | 6.59(1.33)*** | 8.68(.565)*** | 5.99(.448)*** |
| PGS for IQ | 0.89(.456)* | 0.85(1.01) | 3.85(2.16)+ | 2.11(1.05)* | 1.39(.413)*** | 1.47(.385)*** |
| **Population Admixture** |  |  |  |  |  |  |
| PC1 | -37.1 | -1,195.5*** | 204.1 | -485.7+ | -436.7** | -292.8*** |
| PC2 | 330.3 | 214.4 | 329.3 | 457.4** | 320.7*** | 347.1*** |
| PC3 | -370.6+ | -839.2** | -184.6 | -364.1*** | -375.6*** | -406.2*** |
| PC4 | 11.0 | 184.8+ | 667.5** | 53.6 | 31.5 | 46.6+ |
| PC5 | 167.2 | 124.9 | 143.3 | 75.4 | 155.0*** | 173.5*** |
| PC6 | -219.5** | 249.8 | 22.2 | 284.1** | -110.7*** | -95.7*** |
| PC7 | -16.7 | 120.9 | -2.19 | -63.8 | 46.1+ | 35.4 |
| PC8 | -9.8 | -85.1 | -214.8 | 86.4 | -83.0** | -81.0** |
| PC9 | 28.0 | -127.9 | 52.6 | 193.2* | 2.53 | -19.8 |
| PC10 | -2.5 | 107.9 | 200.3 | 57.5 | 13.2 | 25.8 |
| **Constant** | 51.4(1.72)*** | 61.4(2.97)*** | 54.3(6.16)*** | 49.2(1.99)*** | 48.3(.886)*** | 50.3(.279)*** |
| **Random Effects** |  |  |  |  |  |  |
| $\sigma_u^2$, family-level | 2.75(.040)*** | 2.97(.044)*** | 2.89(.147)*** | 2.82(.107)*** | 2.77(.036)*** | 2.83(.029)*** |
| $\sigma_v^2$, person-level | 2.54(.056)*** | 2.29(.138)*** | 2.70(.196)*** | 2.59(.159)*** | 2.57(.050)*** | 2.53(.047)*** |
| $\sigma_e^2$, wave-level | 2.77(.010)*** | 2.79(.018)*** | 2.79(.035)*** | 2.86(.022)*** | 2.79(.009)*** | 2.79(.008)*** |
| **Model-level Parameters** |  |  |  |  |  |  |
| (-2)LogLikelihood | 85814 | 29850 | 7826 | 19908 | 113654 | 143590 |
| Person-observations | 9,441 | 3,253 | 847 | 2,150 | 12,438 | 15,691 |
| Number of Families | 4,374 | 1,510 | 402 | 1,024 | 5,785 | 7,290 |
| Number of Persons | 4,860 | 1,677 | 438 | 1,103 | 6,401 | 8,078 |
| OLS R-squared | 0.044 | 0.049 | 0.148 | 0.106 | 0.114 | 0.154 |

\*\*\* p<0.001, \*\* p<0.01, \* p<0.05, + p<0.1;
Only indicators of level of significance are provided for the estimated PCs.

The models in Table 4 included all SES components as well as the two ability-related PGSs. These models could be compared with the models in Table 2 to assess the impact of the two PGSs on the effects of SES context. All SES components that were statistically significant in the model without the two PGSs remained significant in the model with the PGSs. As expected, the inclusion of the PGSs attenuated the SES effects. In most cases, the inclusion of the PGSs reduced the SES effects in models in Table 2 by about 10% or less. For example, in the white sample in Table 2, the two categories of mother’s education “high-school graduation/some college” and “at least college degree”, respectively, were associated with 6.95 and 13.5 additional points in the verbal ability test score relative to “less than high school” in the model without the PGSs. The two comparable estimates in the model with the two PGSs in Table 4 were 6.26 and 12.3, which represented reductions of 9.9% and 9.2%, respectively. In the black sample, the two comparable reductions were about zero.

**Table 4.**
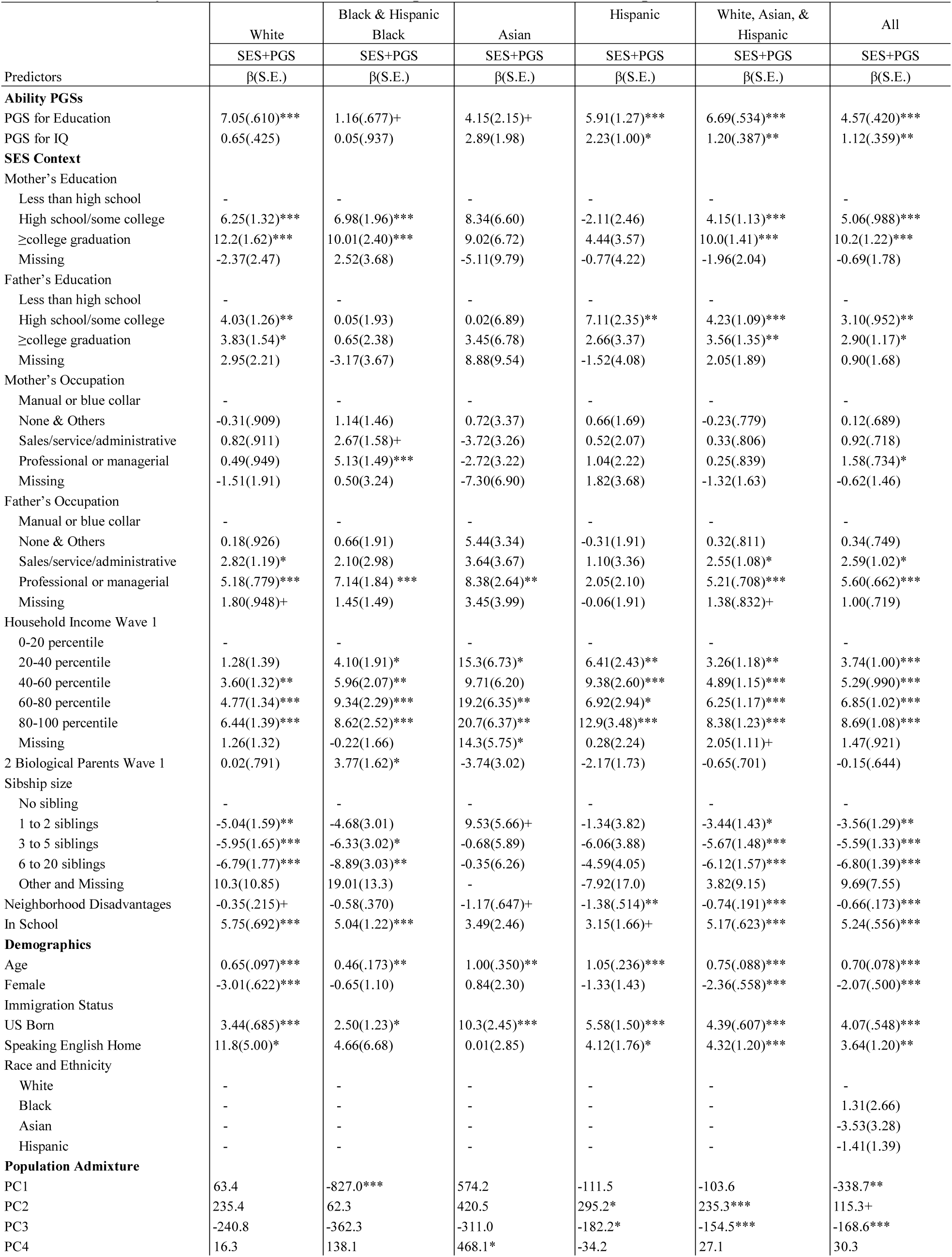
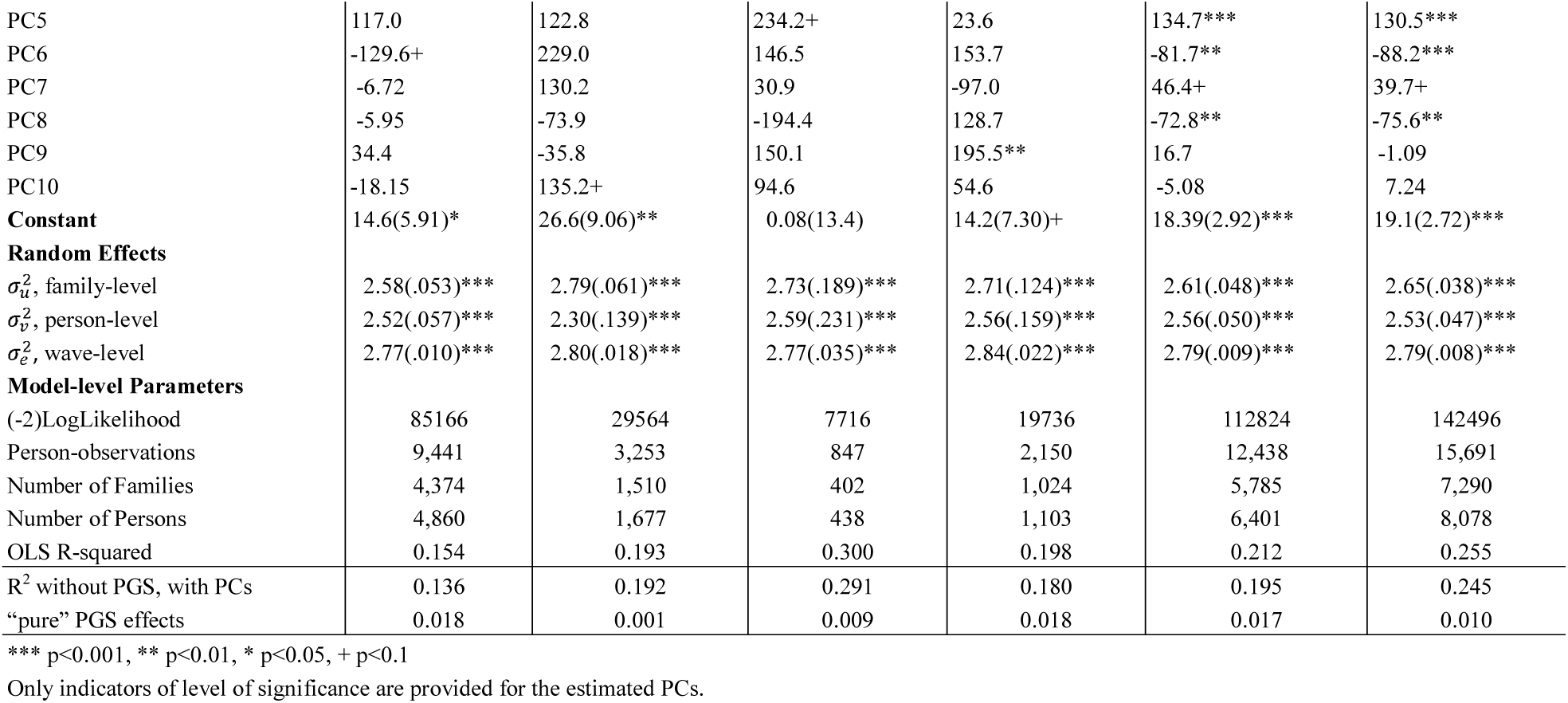
Random-effects models of twice-measured verbal ability: Coefficients (standard errors) of SES context and ability-related PGSs in ethnicities-specific and -combined samples of Add Health

The incremental R^2^s or the amount of R^2^s due to the two PGSs above and beyond the SES components already included in the models represented a measure of the overall genomic influence on verbal ability. They could be obtained by subtracting the R^2^s of the models containing SES context and the principal components from the R^2^s in the models in Table 4. The incremental R^2^s or the R^2^s of “pure” PGS effects were 1.8%, 0.1%, 1%, 1.8%, 1.7%, and 1% for whites, blacks, Asians, Hispanic whites, the combined sample and the overall sample, respectively. These R^2^s were obtained after removing the impact of the principal components.

The models in Table 5 added a set of predictors to models in Table 4 measuring general health and health behaviors and a set of non-ability PGSs. The overall impact of these added predictors on the previous models of SES context and ability-related PGSs was quite modest. The estimates of SES context and the ability PGSs in Table 4 were generally reduced slightly. Most of these added estimates were not statistically significant. A few exceptions included negative effects of marijuana use, smoking and the PGS for conscientiousness in the white sample.

**Table 5.**
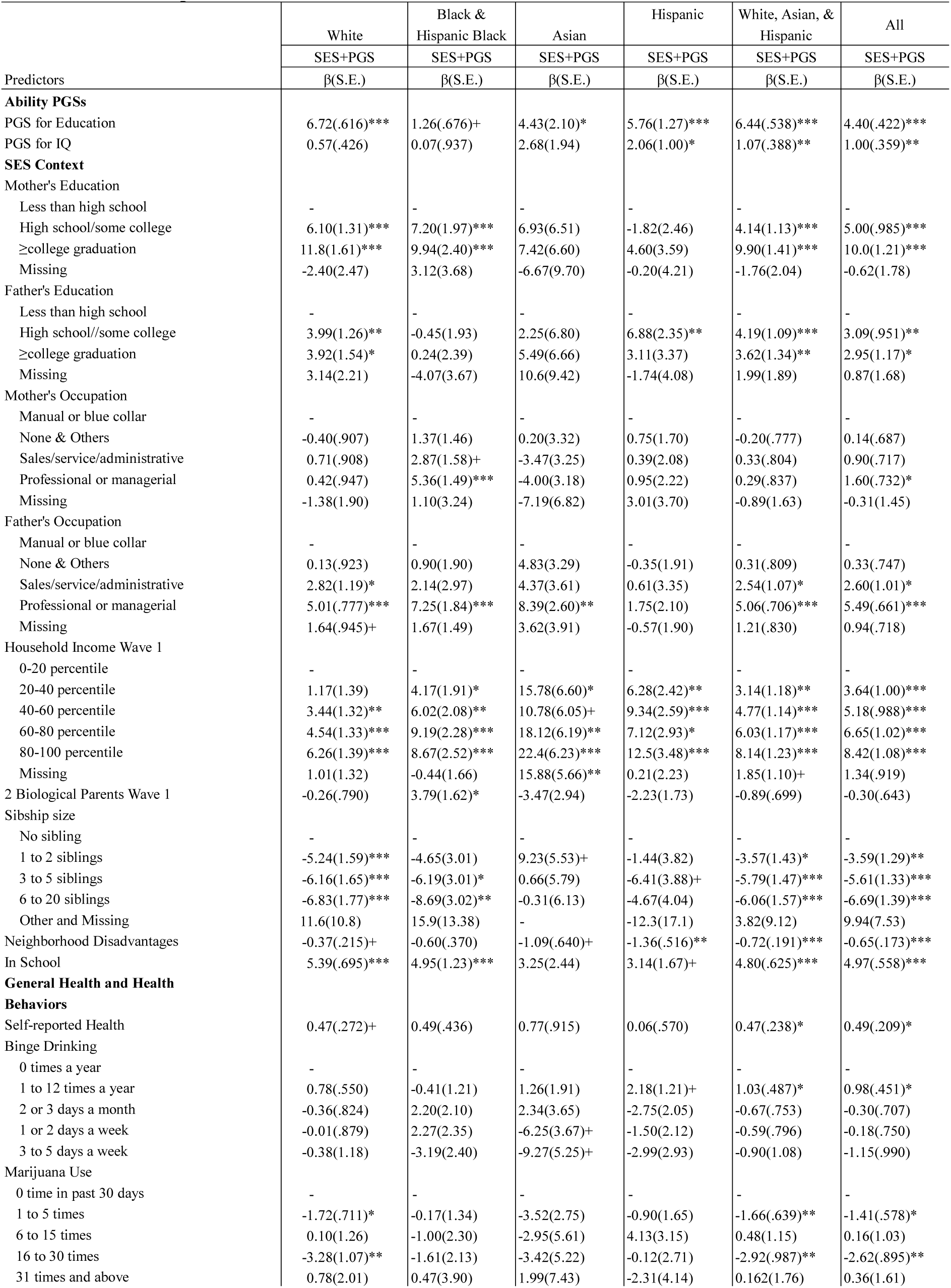
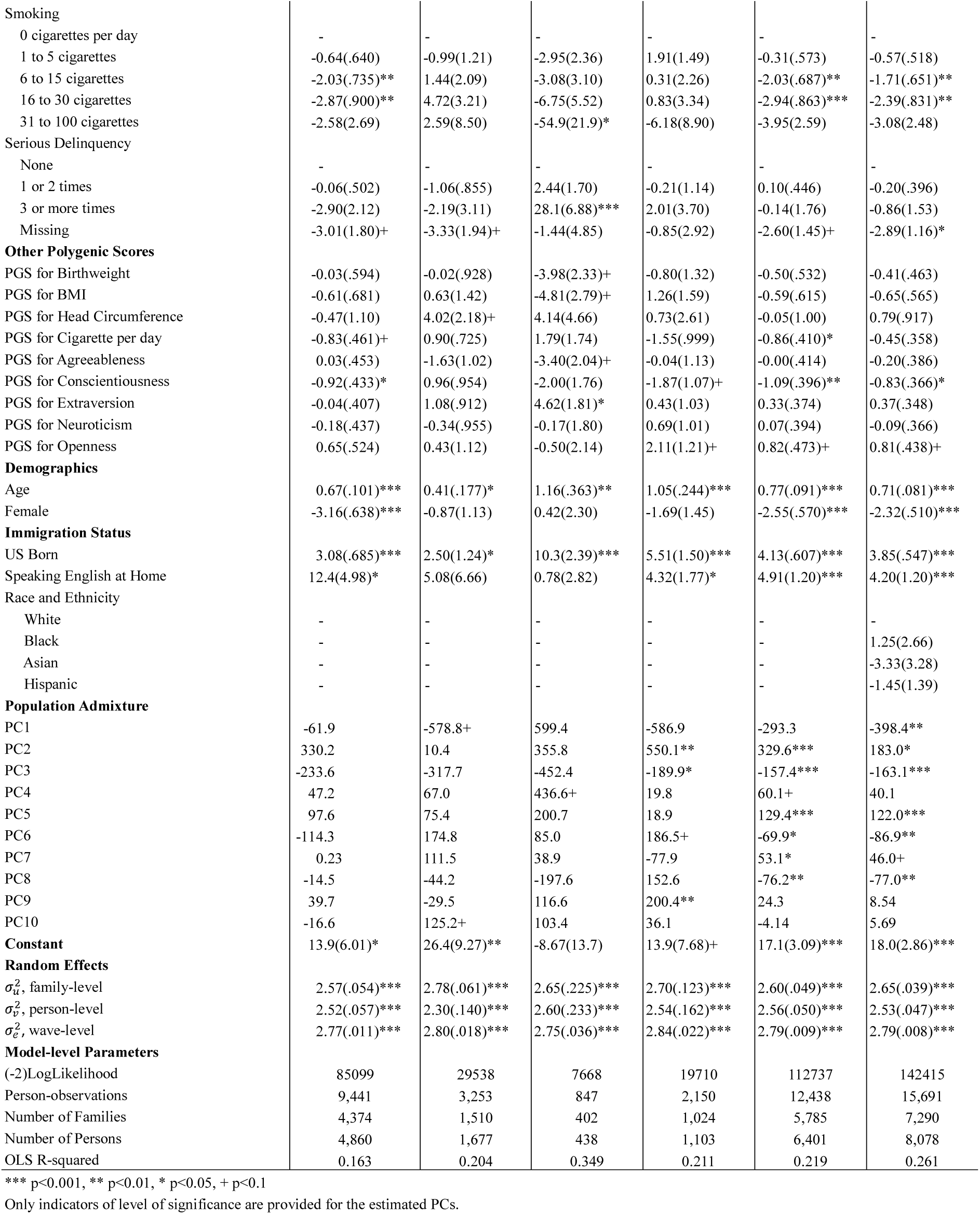
Random-effects models of twice-measured verbal ability: Coefficients (standard errors) of SES context, ability-related PGSs, general health and health behaviors, and other PGSs in ethnicities-specific and -combined samples of Add Health

The models in Table 6 used SES components and the two PGSs to predict the Wave-3 verbal ability while adjusting for the Wave-1 verbal ability measured about seven years before the Wave-3 verbal ability (Equation 2). A total of 37 individuals were excluded because an individual had to contribute two measures of verbal ability to be included in this analysis. Little surprise that the PVT score at Wave 1 was highly predictive of the PVT score at Wave 3. In the white sample, an increase of one point in the Wave-1 PVT score was associated with an increase of about 0.57 points in the Wave-3 PVT score. The effect of one standard deviation of the education PGS was reduced from 7.0 points in Table 4 to 2.2 points in Table 6; this effect was still highly significant. The PGS for intelligence lost its statistical significance in all samples.

Adjusting for the Wave-1 PVT, the effects of SES context were expected to be largely washed out. This was what we observed. Most of the SES components no longer significantly predicted verbal ability at Wave 3. The two remarkable exceptions were years of education by Wave 3 and neighborhood disadvantage at Wave 1. The former was significantly predictive of Wave-3 PVT in every ethnic sample, each additional year of schooling associated with about two points on the PVT score. It was neighborhood disadvantages in earlier life at Wave 1 rather than at Wave 3 that turned out to be negatively associated with verbal ability; both were included in the models. Each standard deviation of neighborhood disadvantages at Wave 1 was negatively associated with about one point in PVT in the white sample. Living with two biological parents was positively associated with an increase of 3.6 points of PVT in the black sample. Its positive effect tended to be washed out or reversed in the other ethnic samples once conditional on Wave-1 PVT.

**Table 6.**
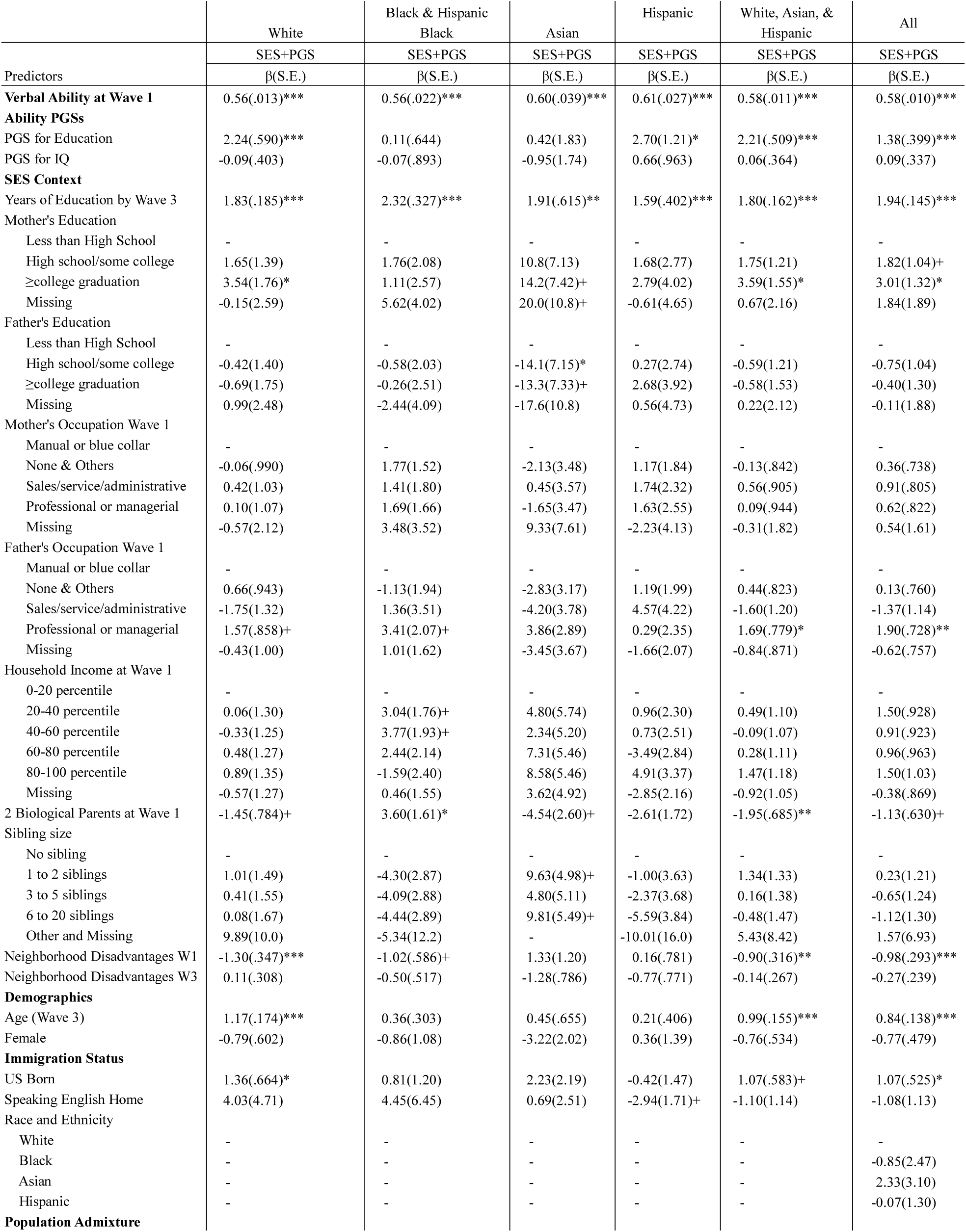
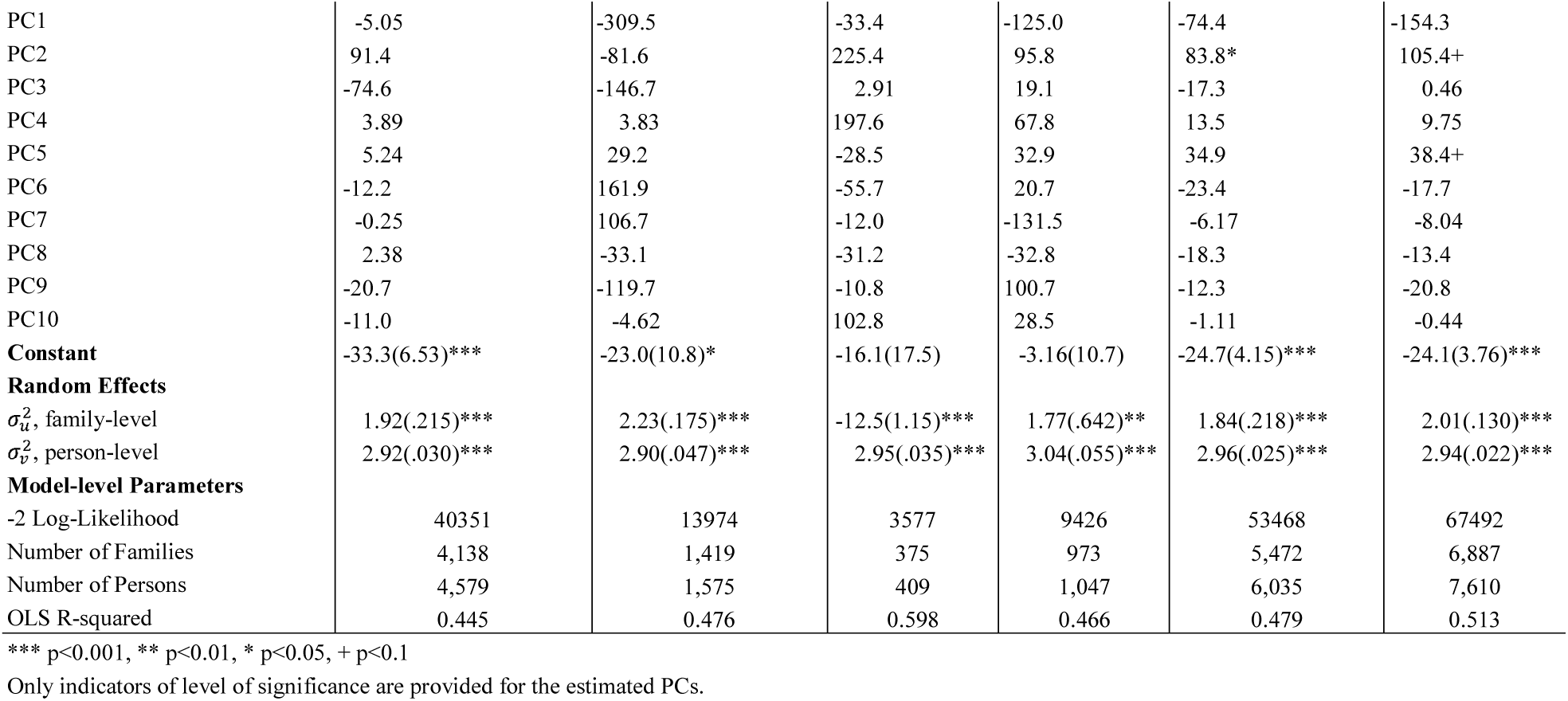
Random-effects models of Wave 3 verbal ability: Coefficients (standard errors) of SES context and ability-related PGSs in ethnicities-specific and -combined samples of Add Health, Conditional on verbal ability at Wave 1

Table 7 showed the main effects and GE interaction effects on twice-measured verbal ability between a PGS composite constructed from the two ability-related PGSs and a SES composite in the black sample, the combined sample, and the overall sample. The effects of the two composites in the main-effects model were significantly predictive of verbal ability in the three analyses. Out of the three GE interaction models, the interaction term was significant at the level of 10% in the black sample and the level of <1% in the overall sample while the term was not significant in the combined sample. In the black sample, those with a value of 1 in the PGS composite and a value of 1 in the SES composite had a PVT score of 60.47 (60.47=45.8+1.9+9.47+3.3) points, which includes 9.47 points of the main effect of the SES composite and 1.9 points of the main effects of the PGS composite, and 3.3 points of additional boost because of GE interaction.

**Table 7.** Random-effects models of twice-measured verbal ability: GE interaction coefficients (standard errors) between a SES composite and a composite of ability-related PGSs in ethnicities-specific and - combined samples of Add Health

| Predictors | Main Effects |  |  | Interaction |  |  |
| --- | --- | --- | --- | --- | --- | --- |
|  | Black & Hispanic Black | White, Asian, & Hispanic | All | Black & Hispanic Black | White, Asian, & Hispanic | All |
|  | SES | SES | SES | SES | SES | SES |
| | $\beta$(S.E.) | $\beta$(S.E.) | $\beta$(S.E.) | $\beta$(S.E.) | $\beta$(S.E.) | $\beta$(S.E.) |
| **Ability PGS composite** | 2.38(1.12)* | 7.47(.616)*** | 6.34(.543)*** | 1.90(1.16) | 7.46(.616)*** | 6.24(.544)*** |
| **SES Composite** | 4.99(1.17)*** | 9.49(.673)*** | 8.50(.585)*** | 9.47(2.83)*** | 9.09(.824)*** | 8.62(.586)*** |
| **PGS x SES** |  |  |  | 3.30(1.90)+ | 1.05(1.25) | 1.69(.616)** |
| **Demographics** |  |  |  |  |  |  |
| Age | -0.08(.092) | 0.21(.046)*** | 0.15(.041)*** | -0.08(.092) | 0.21(.046)*** | 0.15(.041)*** |
| Female | -2.18(1.17)+ | -2.64(.586)*** | -2.59(.527)*** | -2.20(1.17)+ | -2.64(.586)*** | -2.62(.527)*** |
| 2 Biological Parents W1 | 9.28(1.36)*** | 2.92(.602)*** | 3.95(.553)*** | 9.27(1.36)*** | 2.91(.602)*** | 3.93(.553)*** |
| **Immigration Status** |  |  |  |  |  |  |
| US Born | 3.78(1.32)** | 4.40(.639)*** | 4.33(.578)*** | 3.83(1.32)** | 4.41(.639)*** | 4.36(.578)*** |
| Speaking English Home | 8.78(7.39) | 8.18(1.24)*** | 7.60(1.25)*** | 8.39(7.39) | 8.20(1.24)*** | 7.54(1.25)*** |
| **Race and Ethnicity** |  |  |  |  |  |  |
| White | - | - | - | - | - | - |
| Black | - | - | 2.52(2.81) | - | - | 2.85(2.81) |
| Asian | - | - | -1.60(3.48) | - | - | -1.53(3.48) |
| Hispanic | - | - | -1.99(1.47) | - | - | -1.93(1.47) |
| **Population Admixture** |  |  |  |  |  |  |
| PC1 | -1,042.2*** | -326.3* | -486.8*** | -1,024.2*** | -329.2* | -492.3*** |
| PC2 | 238.0 | 171.1*** | 110.3 | 212.4 | 169.7*** | 111.2+ |
| PC3 | -690.7* | -277.5*** | -258.7*** | -676.7* | -278.6*** | -258.2*** |
| PC4 | 170.4 | 37.4 | 35.2 | 174.4+ | 37.1 | 35.6 |
| PC5 | 96.4 | 149.7*** | 140.6*** | 95.1 | 149.6*** | 140.7*** |
| PC6 | 253.4 | -132.1*** | -139.2*** | 258.4 | -132.1*** | -138.4*** |
| PC7 | 65.4 | 32.0 | 22.3 | 68.9 | 32.1 | 22.7 |
| PC8 | -59.1 | -73.2** | -78.5** | -59.8 | -73.3** | -78.1** |
| PC9 | -90.3 | -8.34 | -24.7 | -91.8 | -8.53 | -24.4 |
| PC10 | 135.1+ | -0.97 | 14.0 | 134.3+ | -0.88 | 13.6 |
| **Constant** | 46.2(8.08)*** | 35.6(1.68)*** | 36.6(1.70)*** | 45.7(8.08)*** | 35.5(1.68)*** | 36.7(1.70)*** |
| **Random Effects** |  |  |  |  |  |  |
| $\sigma_u^2$, family-level | 2.92(.049)*** | 2.69(.042)*** | 2.74(.034)*** | 2.92(.049)*** | 2.69(.043)*** | 2.74(.034)*** |
| $\sigma_v^2$, person-level | 2.29(.143)*** | 2.59(.049)*** | 2.55(.046)*** | 2.29(.143)*** | 2.59(.049)*** | 2.55(.046)*** |
| $\sigma_e^2$, wave-level | 2.79(.018)*** | 2.79(.009)*** | 2.79(.008)*** | 2.79(.018)*** | 2.79(.009)*** | 2.79(.008)*** |
| **Model-level Parameters** |  |  |  |  |  |  |
| (-2)LogLikelihood | 28837 | 110832 | 139734 | 28834 | 110831 | 139726 |
| Person-observations | 3,153 | 12,163 | 15,316 | 3,153 | 12,163 | 15,316 |
| Number of Families | 1,460 | 5,642 | 7,097 | 1,460 | 5,642 | 7,097 |
| Number of Persons | 1,619 | 6,235 | 7,854 | 1,619 | 6,235 | 7,854 |
| OLS R-squared | 0.098 | 0.151 | 0.193 | 0.100 | 0.151 | 0.194 |

\*\*\* p<0.001, \*\* p<0.01, \* p<0.05, + p<0.1
Only indicators of level of significance are provided for the estimated PCs.

## DISCUSSION AND CONCLUSION

The models of SES context predicted verbal ability well. After demographic variables and immigration status were adjusted, mother’s education, father’s education, mother’s occupation, father’s occupation, household income, sibship size, neighborhood disadvantage, and in school over the past year were all significantly and largely independently associated with verbal ability. Important differences across ethnic groups did exist. In the black sample, father’s education was much less influential and mother’s occupation was much more influential than their counterparts in the white sample. The differences could be explained by the much higher proportion of female-headed families in the black sample. Table 1 indicated that 30% of black respondents vs 50% of white ones reported living with 2-biological parents at Wave 1. The estimates in the Asian sample tended to be less statistically significant for at least three reasons. Being only one tenth of the white sample, the Asian sample size was likely to be a major cause of the non-significance. The Asian sample in Add Health was highly heterogeneous with respect to the age at which they immigrated to the United States, which was a crucial factor for verbal ability in English. The cultural background in the Asian sample was also highly heterogeneous. The Asian sample consisted of the Chinese, Filipinos, Japanese, Koreans, and Vietnamese. These patterns of cross-ethnicity differences largely persisted when the ability PGSs were included in the models.

In the combined sample of whites, Asians, and Hispanic whites, when the ability PGSs and SES context were in a model simultaneously (Table 4), both tended to continue to significantly predict verbal ability. All SES predictors that were predictive of verbal ability without the PGSs remained significantly predictive of verbal ability with the PGSs. The model that included both SES factors and the ability PGSs yielded an *R^2^* of 21.2%, which represented an increase of an *R^2^* of 1.7% over the model of SES context, excluding the impact the PCs. The effect of the education PGS and the effect of the intelligence PGS were reduced about 23% and 28%, respectively, when SES context were added. The reductions in the effects of SES context were mostly about 10% or less.

The overall payoff of adding general health and health behaviors and non-ability PGSs is negligible. Only smoking and marijuana use were negatively associated with verbal ability and these effects were small. Adding the non-ability PGSs did not yield any additional significant explanatory power to the model. More importantly for the purpose of this analysis, adding these two classes of factors to the model did not affect the estimates of SES context and the effects of the two ability PGSs already in the model. In contrast to SES predictors and the ability PGSs, these other factors were of less consequences to verbal ability.

Moving Wave-1 verbal ability from the left side of the equation to the right side or conditioning the analysis of Wave-3 verbal ability on Wave-1 verbal ability further tested the importance of SES context. A coefficient of about 0.6 of Wave-1 verbal ability indicated that Wave-1 verbal ability predicted Wave-3 verbal ability with 60% accuracy. With the Wave-1 verbal ability adjusted, the estimated effects of the PGSs and SES predictors carried a distinct meaning. These effects represented the effects above and beyond those that had gone into Wave-1 verbal ability, or above and beyond the effects of the measured and unmeasured PGSs, SES and other predictors that had already acted upon Wave-1 verbal ability. The PGS for education was still significantly predictive of Wave-3 verbal ability, but the effect size was reduced by about two thirds. Two SES predictors stood out: number of years of schooling by Wave 3 and neighborhood disadvantage at Wave 1. The large effect of year of schooling by Wave 3, which was significant in every ethnic sample, was particularly noteworthy. The coefficient of 1.83 implied that an additional year of schooling was associated with almost two points of verbal ability. Assessing causal effects of schooling on verbal ability was difficult. While more schooling helps grow verbal ability, individuals with higher verbal ability are likely to seek and attain more education. We took three measures to address the difficulty. First, the model controlled for an earlier version of verbal ability, which was equivalent to controlling for the ability for seeking and attaining education. Second, the model controlled for age at which Wave-3 test was taken since age was likely to be positively correlated with the amount of schooling. Third, we measured schooling by the number of years of schooling before verbal ability was taken at Wave 3.

Our gene-environment interaction analysis was carried out by interacting a composite PGS based on the two ability-related PGSs and a composite score for SES context. Two of the three ethnic samples tested confirmed the theoretical prediction that favorable SES context boosted the level of cognitive ability predicted by the ability PGSs, augmenting evidence for the important role of SES context for cognition. The positive effect of GE interaction in the black sample was at least twice as large as the counterpart in the non-black sample. This contrasted the much smaller main effect of the education PGS in the black sample than in the non-black sample.

Carrying out genomic analysis by ethnic group led to a number of insights. The effects of ability PGSs and SES appeared broadly similar across whites, Asians and non-Hispanic whites. Although the effect of the education PGS tended to predict verbal ability in the black sample, it was much smaller in size and less significant. We already discussed the lack of African participation in the current GWAS, which resulted in significantly less than full coverage of African genetic variants related to cognitive ability. The much smaller black sample was likely part of the explanation also. Lastly, the much larger positive GE interaction effect in the black sample indicated that unfavorable SES context played a much larger role in holding back black youths from realizing their genomic potential for cognitive development. A fuller understanding of how ability PGSs are related to cognitive development in the black sample has to wait for a significant expansion of GWAS into the African populations.

The findings from models that included both genomic and SES measures require careful interpretation. Parental genomes are a common origin of offspring’s genome and SES context. Parental genomes transmit genetic materials to offspring and shape parental SES such as parental education, occupation, and income as well as the neighborhood the family lives in and the schools the children attend. This common origin is the reason for the correlation between SES context and children’s ability PGSs. A recent Icelandic study of genomic analysis of educational attainment separated the parental alleles that have been transmitted to the offspring and the parental alleles that have not been transmitted to the offspring (Kong et al. 2018). Although two generations of genomic data are unavailable in Add Health, the approach used in the study suggests a way of estimating the extent to which SES context is genomic.

The inclusion of the two ability PGSs and SES context in the same model yielded a number of invaluable findings. It established the importance of genomic roots of verbal ability. The estimated effect of the PGS for education was substantial in size, highly significant, and largely invariant to whether the model included general health and health behaviors, and non-ability PGSs. The ability PGSs accounted for 1.7% of the incremental R^2^ of verbal ability in the combined sample. In all probability, the measurement of genomic root of verbal ability will improve. Substantially larger samples than currently used could be assembled to hunt more genes in fresh GWAS, which focus on common alleles with MAF greater than 5%. A recent study (Marouli et al. 2017) showed that SNPs with MAF<5% can be examined when an extremely large sample is available and that these rare variants tend to have much larger effects than common variants. These foreseeable advances plus unforeseeable advances will in all likelihood raise the percentage of variance that can be explained by genomic measures.

When both SES context and the ability PGSs were included in a model, the effects of both were estimated with greater accuracy. The estimated effects of SES context were less “contaminated” by parental genomic sources and they were more the effects of non-genomic SES context. The effects of the ability PGSs became more truly a reflection of children’s own genome rather than his or her parents’ genomes. Genetic effects estimated from GWAS measures without adjusting for family environment are generally exaggerated because such estimates count the effects of family genomes as the effects of the offspring’s genome (Kong et al. 2018). In our analysis, adding SES context reduced the effect of the education PGS by more than 20%, which was our estimated amount of exaggeration when the effect of offspring’s education PGS was assessed without adjusting for SES context.

Traditional models of SES context such as those in Table 2 were not “pure” social-science models. With the ability PGSs added to traditional social science models, the relative importance of non-genomic SES context vs. genomic SES context may be assessed. SES context could be viewed as consisting two components: the component due to parental genomics and the component not due to parental genomes. Of the parental genomes that influence verbal ability through nurturing environment, about 50% of the alleles were transmitted to the offspring and the other 50% were not transmitted. The transmitted alleles shared between the parents and the offspring were the reasons why the effects of SES context were reduced by 10% or less. Doubling this 10% or less would be our estimated proportion of the total effects of SES context that was due to parental genomes. The doubling was needed because only 50% of the influential alleles were transmitted.

Given these estimates, SES context remained a dominant force on verbal ability. Non-genomic effects accounted for >80% of SES effects. Even this <20% genomic SES effects could be viewed as environmentally nurturing because they acted from outside the offspring and constituted part of the offspring’s environment. The dominance of SES context was also indicated by the contrast between the R^2^ of 17.5% in the model of SES text (Table 2) and the incremental R^2^ of 1.7%, which was due to the ability PGSs.

While it is true that the current genomic measures of cognitive ability are underestimated, the same could be said about SES context. Presently, SES is measured by crude categories of parental characteristics such education and occupation. Within each of these parental categories, the differences made by idiosyncratic nurturing styles of parents are largely uncaptured.

Social scientists have long considered SES context fundamentally important in shaping life outcomes including cognitive ability (e.g., Duncan, Brooksgunn and Klebanov 1994; Fischer et al. 1996; Hauser 2010; McLeod and Shanahan 1993). With genomic influences measured at individual level and included in the model, we confirmed the importance of SES context. After all, the worth of cognitive ability including verbal ability depends heavily on the context of modern education and society. At the same time, the modern educational system and society depend on social fabrics such as family, schools, and neighborhoods to bring about the intellectual potential of individuals. The eminence of these social fabrics will persist unless their functions are replaced by institutions other than family, schools, and neighborhoods.

---

**Appendix 1.**
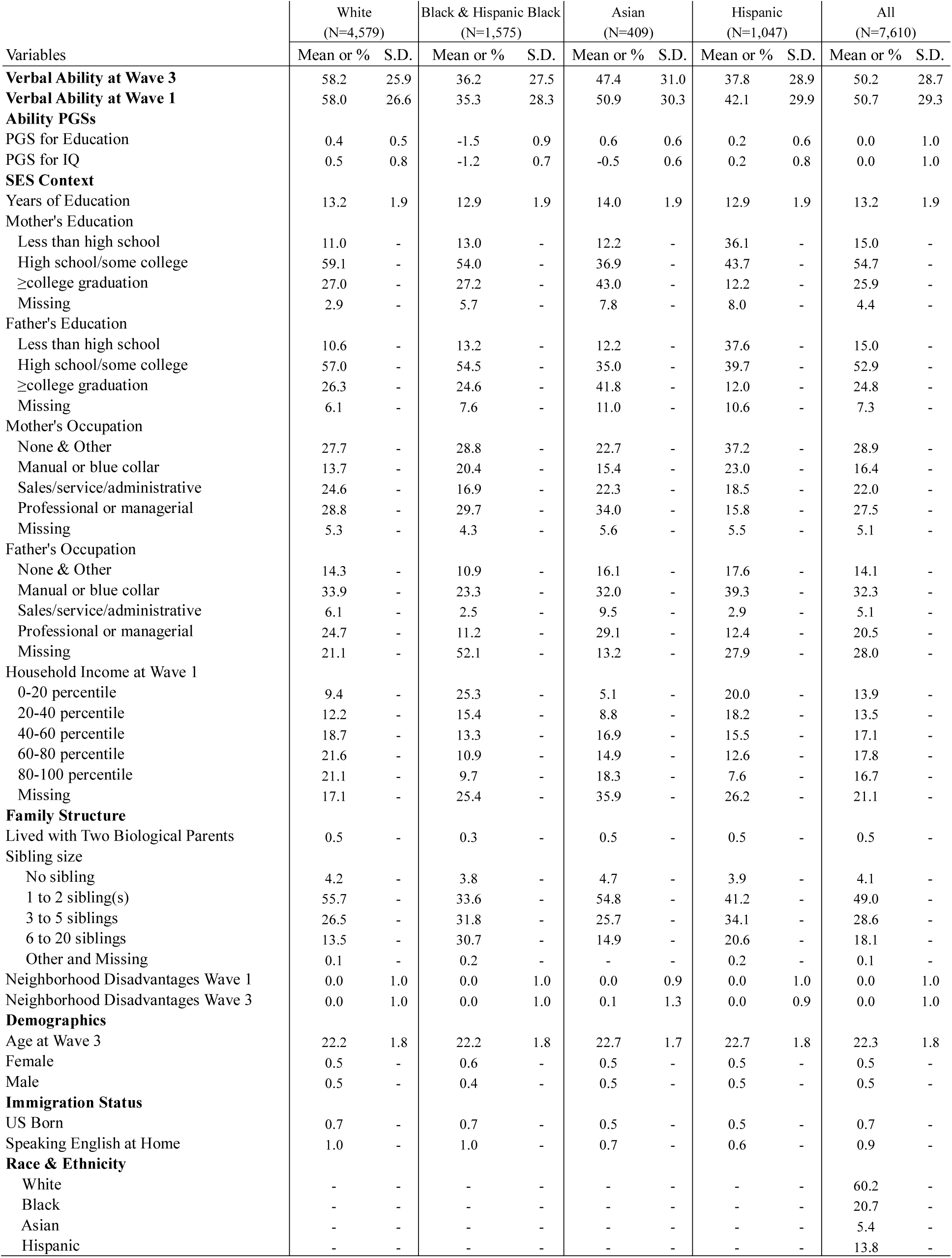
Descriptive statistics for the samples used in the analysis conditional on Wave-1 verbal ability

| Variables | White<br>(N=4,579) |  | Black & Hispanic Black<br>(N=1,575) |  | Asian<br>(N=409) |  | Hispanic<br>(N=1,047) |  | All<br>(N=7,610) |  |
| --- | --- | --- | --- | --- | --- | --- | --- | --- | --- | --- |
|  | Mean or % | S.D. | Mean or % | S.D. | Mean or % | S.D. | Mean or % | S.D. | Mean or % | S.D. |
| **Verbal Ability at Wave 3** | 58.2 | 25.9 | 36.2 | 27.5 | 47.4 | 31.0 | 37.8 | 28.9 | 50.2 | 28.7 |
| **Verbal Ability at Wave 1** | 58.0 | 26.6 | 35.3 | 28.3 | 50.9 | 30.3 | 42.1 | 29.9 | 50.7 | 29.3 |
| **Ability PGSs** |  |  |  |  |  |  |  |  |  |  |
| PGS for Education | 0.4 | 0.5 | -1.5 | 0.9 | 0.6 | 0.6 | 0.2 | 0.6 | 0.0 | 1.0 |
| PGS for IQ | 0.5 | 0.8 | -1.2 | 0.7 | -0.5 | 0.6 | 0.2 | 0.8 | 0.0 | 1.0 |
| **SES Context** |  |  |  |  |  |  |  |  |  |  |
| Years of Education | 13.2 | 1.9 | 12.9 | 1.9 | 14.0 | 1.9 | 12.9 | 1.9 | 13.2 | 1.9 |
| Mother's Education |  |  |  |  |  |  |  |  |  |  |
| Less than high school | 11.0 | - | 13.0 | - | 12.2 | - | 36.1 | - | 15.0 | - |
| High school/some college | 59.1 | - | 54.0 | - | 36.9 | - | 43.7 | - | 54.7 | - |
| ≥college graduation | 27.0 | - | 27.2 | - | 43.0 | - | 12.2 | - | 25.9 | - |
| Missing | 2.9 | - | 5.7 | - | 7.8 | - | 8.0 | - | 4.4 | - |
| Father's Education |  |  |  |  |  |  |  |  |  |  |
| Less than high school | 10.6 | - | 13.2 | - | 12.2 | - | 37.6 | - | 15.0 | - |
| High school/some college | 57.0 | - | 54.5 | - | 35.0 | - | 39.7 | - | 52.9 | - |
| ≥college graduation | 26.3 | - | 24.6 | - | 41.8 | - | 12.0 | - | 24.8 | - |
| Missing | 6.1 | - | 7.6 | - | 11.0 | - | 10.6 | - | 7.3 | - |
| Mother's Occupation |  |  |  |  |  |  |  |  |  |  |
| None & Other | 27.7 | - | 28.8 | - | 22.7 | - | 37.2 | - | 28.9 | - |
| Manual or blue collar | 13.7 | - | 20.4 | - | 15.4 | - | 23.0 | - | 16.4 | - |
| Sales/service/administrative | 24.6 | - | 16.9 | - | 22.3 | - | 18.5 | - | 22.0 | - |
| Professional or managerial | 28.8 | - | 29.7 | - | 34.0 | - | 15.8 | - | 27.5 | - |
| Missing | 5.3 | - | 4.3 | - | 5.6 | - | 5.5 | - | 5.1 | - |
| Father's Occupation |  |  |  |  |  |  |  |  |  |  |
| None & Other | 14.3 | - | 10.9 | - | 16.1 | - | 17.6 | - | 14.1 | - |
| Manual or blue collar | 33.9 | - | 23.3 | - | 32.0 | - | 39.3 | - | 32.3 | - |
| Sales/service/administrative | 6.1 | - | 2.5 | - | 9.5 | - | 2.9 | - | 5.1 | - |
| Professional or managerial | 24.7 | - | 11.2 | - | 29.1 | - | 12.4 | - | 20.5 | - |
| Missing | 21.1 | - | 52.1 | - | 13.2 | - | 27.9 | - | 28.0 | - |
| Household Income at Wave 1 |  |  |  |  |  |  |  |  |  |  |
| 0-20 percentile | 9.4 | - | 25.3 | - | 5.1 | - | 20.0 | - | 13.9 | - |
| 20-40 percentile | 12.2 | - | 15.4 | - | 8.8 | - | 18.2 | - | 13.5 | - |
| 40-60 percentile | 18.7 | - | 13.3 | - | 16.9 | - | 15.5 | - | 17.1 | - |
| 60-80 percentile | 21.6 | - | 10.9 | - | 14.9 | - | 12.6 | - | 17.8 | - |
| 80-100 percentile | 21.1 | - | 9.7 | - | 18.3 | - | 7.6 | - | 16.7 | - |
| Missing | 17.1 | - | 25.4 | - | 35.9 | - | 26.2 | - | 21.1 | - |
| **Family Structure** |  |  |  |  |  |  |  |  |  |  |
| Lived with Two Biological Parents | 0.5 | - | 0.3 | - | 0.5 | - | 0.5 | - | 0.5 | - |
| Sibling size |  |  |  |  |  |  |  |  |  |  |
| No sibling | 4.2 | - | 3.8 | - | 4.7 | - | 3.9 | - | 4.1 | - |
| 1 to 2 sibling(s) | 55.7 | - | 33.6 | - | 54.8 | - | 41.2 | - | 49.0 | - |
| 3 to 5 siblings | 26.5 | - | 31.8 | - | 25.7 | - | 34.1 | - | 28.6 | - |
| 6 to 20 siblings | 13.5 | - | 30.7 | - | 14.9 | - | 20.6 | - | 18.1 | - |
| Other and Missing | 0.1 | - | 0.2 | - | - | - | 0.2 | - | 0.1 | - |
| Neighborhood Disadvantages Wave 1 | 0.0 | 1.0 | 0.0 | 1.0 | 0.0 | 0.9 | 0.0 | 1.0 | 0.0 | 1.0 |
| Neighborhood Disadvantages Wave 3 | 0.0 | 1.0 | 0.0 | 1.0 | 0.1 | 1.3 | 0.0 | 0.9 | 0.0 | 1.0 |
| **Demographics** |  |  |  |  |  |  |  |  |  |  |
| Age at Wave 3 | 22.2 | 1.8 | 22.2 | 1.8 | 22.7 | 1.7 | 22.7 | 1.8 | 22.3 | 1.8 |
| Female | 0.5 | - | 0.6 | - | 0.5 | - | 0.5 | - | 0.5 | - |
| Male | 0.5 | - | 0.4 | - | 0.5 | - | 0.5 | - | 0.5 | - |
| **Immigration Status** |  |  |  |  |  |  |  |  |  |  |
| US Born | 0.7 | - | 0.7 | - | 0.5 | - | 0.5 | - | 0.7 | - |
| Speaking English at Home | 1.0 | - | 1.0 | - | 0.7 | - | 0.6 | - | 0.9 | - |
| **Race & Ethnicity** |  |  |  |  |  |  |  |  |  |  |
| White | - | - | - | - | - | - | - | - | 60.2 | - |
| Black | - | - | - | - | - | - | - | - | 20.7 | - |
| Asian | - | - | - | - | - | - | - | - | 5.4 | - |
| Hispanic | - | - | - | - | - | - | - | - | 13.8 | - |

**Appendix 2.** Descriptive statistics for GE Interaction Analysis: Add Health by Ethnicity

| Variable | Black & Hispanic Black<br>(N=1,619) |  | White, Asian, & Hispanic<br>(N=6,235) |  | All<br>(N=7,854) |  |
| --- | --- | --- | --- | --- | --- | --- |
|  | Mean or % | S.D. | Mean or % | S.D. | Mean or % | S.D. |
| **Verbal Ability** | 36.1 | 28.0 | 54.6 | 28.0 | 50.8 | 28.9 |
| **Ability PGS Composite** | -1.3 | 0.6 | 0.4 | 0.5 | 0.0 | 0.9 |
| **SES Composite** | 0.2 | 0.4 | 0.0 | 0.4 | 0.0 | 0.4 |
| **Demographics** |  |  |  |  |  |  |
| Age | 19.1 | 3.6 | 19.2 | 3.6 | 19.2 | 3.6 |
| Gender |  |  |  |  |  |  |
| Female | 0.6 | - | 0.5 | - | 0.5 | - |
| Male | 0.4 | - | 0.5 | - | 0.5 | - |
| 2 Biological Parents | 0.3 | - | 0.5 | - | 0.5 | - |
| Immigration Status |  |  |  |  |  |  |
| US Born | 0.7 | - | 0.7 | - | 0.7 | - |
| Speaking English at Home | 1.0 | - | 0.9 | - | 0.9 | - |
| Race & Ethnicity |  |  |  |  |  |  |
| White | - | - | - | - | 60.6 | - |
| Black | - | - | - | - | 20.6 | - |
| Asian | - | - | - | - | 5.3 | - |
| Hispanic | - | - | - | - | 13.5 | - |

## Acknowledgements

This research uses data from Add Health, a program project designed by J. Richard Udry, Peter S. Bearman, and Kathleen Mullan Harris, and funded by a grant P01-HD31921 from the National Institute of Child Health and Human Development, with cooperative funding from 17 other agencies (http://www.cpc.unc.edu/addhealth/contract.html).

## THE REFERENCES

Allen, H. L., K. Estrada, G. Lettre, S. I. Berndt, M. N. Weedon, F. Rivadeneira, C. J. Willer, A. U. Jackson, S. Vedantam, S. Raychaudhuri, T. Ferreira, A. R. Wood, R. J. Weyant, A. V. Segre, E. K. Speliotes, E. Wheeler, N. Soranzo, J. H. Park, J. Yang, D. Gudbjartsson, N. L. Heard-Costa, J. C. Randall, L. Qi, A. V. Smith, R. Magi, T. Pastinen, L. Liang, I. M. Heid, J. Luan, G. Thorleifsson, T. W. Winkler, M. E. Goddard, K. S. Lo, C. Palmer, T. Workalemahu, Y. S. Aulchenko, A. Johansson, M. C. Zillikens, M. F. Feitosa, T. Esko, T. Johnson, S. Ketkar, P. Kraft, M. Mangino, I. Prokopenko, D. Absher, E. Albrecht, F. Ernst, N. L. Glazer, C. Hayward, J. J. Hottenga, K. B. Jacobs, J. W. Knowles, Z. Kutalik, K. L. Monda, O. Polasek, M. Preuss, N. W. Rayner, N. R. Robertson, V. Steinthorsdottir, J. P. Tyrer, B. F. Voight, F. Wiklund, J. F. Xu, J. H. Zhao, D. R. Nyholt, N. Pellikka, M. Perola, J. R. B. Perry, I. Surakka, M. L. Tammesoo, E. L. Altmaier, N. Amin, T. Aspelund, T. Bhangale, G. Boucher, D. I. Chasman, C. Chen, L. Coin, M. N. Cooper, A. L. Dixon, Q. Gibson, E. Grundberg, K. Hao, M. J. Junttila, L. M. Kaplan, J. Kettunen, I. R. Konig, T. Kwan, R. W. Lawrence, D. F. Levinson, M. Lorentzon, B. McKnight, A. P. Morris, M. Muller, J. S. Ngwa, S. Purcell, S. Rafelt, R. M. Salem, E. Salvi, et al. 2010. “Hundreds of variants clustered in genomic loci and biological pathways affect human height.” Nature 467(7317):832–38.

Altshuler, D., L. D. Brooks, A. Chakravarti, F. S. Collins, M. J. Daly, and P. Donnelly. 2005. “A haplotype map of the human genome.” Nature 437(7063):1299–320.

Altshuler, David, Richard M. Durbin, Goncalo R. Abecasis, David R. Bentley, Aravinda Chakravarti, Andrew G. Clark, Francis S. Collins, Francisco M. De la Vega, Peter Donnelly, Michael Egholm, Paul Flicek, Stacey B. Gabriel, Richard A. Gibbs, Bartha M. Knoppers, Eric S. Lander, Hans Lehrach, Elaine R. Mardis, Gil A. McVean, DebbieA Nickerson, Leena Peltonen, Alan J. Schafer, Stephen T. Sherry, Jun Wang, Richard K. Wilson, Richard A. Gibbs, David Deiros, Mike Metzker, Donna Muzny, Jeff Reid, David Wheeler, Jun Wang, Jingxiang Li, Min Jian, Guoqing Li, Ruiqiang Li, Huiqing Liang, Geng Tian, Bo Wang, Jian Wang, Wei Wang, Huanming Yang, Xiuqing Zhang, Huisong Zheng, Eric S. Lander, David L. Altshuler, Lauren Ambrogio, Toby Bloom, Kristian Cibulskis, Tim J. Fennell, Stacey B. Gabriel, David B. Jaffe, Erica Shefler, Carrie L. Sougnez, David R. Bentley, Niall Gormley, Sean Humphray, Zoya Kingsbury, Paula Koko-Gonzales, Jennifer Stone, Kevin J. McKernan, Gina L. Costa, Jeffry K. Ichikawa, Clarence C. Lee, Ralf Sudbrak, Hans Lehrach, Tatiana A. Borodina, Andreas Dahl, Alexey N. Davydov, Peter Marquardt, Florian Mertes, Wilfiried Nietfeld, Philip Rosenstiel, Stefan Schreiber, Aleksey V. Soldatov, Bernd Timmermann, Marius Tolzmann, Michael Egholm, Jason Affourtit, Dana Ashworth, Said Attiya, Melissa Bachorski, Eli Buglione, Adam Burke, Amanda Caprio, Christopher Celone, Shauna Clark, David Conners, Brian Desany, Lisa Gu, Lorri Guccione, Kalvin Kao, Andrew Kebbel, Jennifer Knowlton, Matthew Labrecque, Louise McDade, Craig Mealmaker, Melissa Minderman, Anne Nawrocki, Faheem Niazi, Kristen Pareja, et al. 2010. “A map of human genome variation from population-scale sequencing.” Nature 467(7319):1061–73.

Bell, N. L., K. S. Lassiter, T. D. Matthews, and M. B. Hutchinson. 2001. “Comparison of the Peabody Picture Vocabulary Test-Third Edition and Wechsler Adult Intelligence Scale-Third Edition with university students.” Journal of Clinical Psychology 57(3):417–22.

Beres, Kristee A., Alan S. Kaufman, and Mitchel D. Perlman. 1999. “Assessment of Child Intelligence.” Pp. 65–96 in Handbook of Psychological Assessment edited by Gerald Goldstein and Michel Hersen.

Bouchard, T. J. 1998. “Genetic and environmental influences on adult intelligence and special mental abilities.” Human Biology 70(2):257–79.

Bouchard, T. J., and M. McGue. 1981. “FAMILIAL STUDIES OF INTELLIGENCE - A REVIEW.” Science 212(4498):1055–59.

Braveman, P. A., C. Cubbin, S. Egerter, S. Chideya, K. S. Marchi, M. Metzler, and S. Posner. 2005. “Socioeconomic status in health research - One size does not fit all.” Jama-Journal of the American Medical Association 294(22):2879–88.

Bulik-Sullivan, B. K., P. R. Loh, H. K. Finucane, S. Ripke, J. Yang, N. Patterson, M. J. Daly, A. L. Price, B. M. Neale, and Grp Schizophrenia Working. 2015. “LD Score regression distinguishes confounding from polygenicity in genome-wide association studies.” Nature Genetics 47(3):291-+.

Burt, C., E. Jones, E. Miller, and W. Moodie. 1934. How the mind works. New York: Appleton-Century-Crofts.

Cahan, S., and N. Cohen. 1989. “AGE VERSUS SCHOOLING EFFECTS ON INTELLIGENCE DEVELOPMENT.” Child Development 60(5):1239–49.

Campbell, Jonathan M., and Aila K. Dommestrup. 2010. “Peabody Picture Vocabulary Test.” in Corsini Encyclopedia of Psychology. New York: John Wiley & Sons, Inc.

Carvajal, H., J. E. Hayes, H. R. Miller, D. A. Wiebe, and K. A. Weaver. 1993. “COMPARISONS OF THE VOCABULARY SCORES AND IQS ON THE WECHSLER-INTELLIGENCE-SCALE-FOR-CHILDREN-III AND THE PEABODY-PICTURE-VOCABULARY-TEST-REVISED.” Perceptual and Motor Skills 76(1):28–30.

Cavalli-Sforza, L. Luca, Paolo Menozzi, and Alberto Piazza. 1996. The History and Geography of Human Genes. Princeton, N.J.: Princeton University Press.

Chabris, C. F., B. M. Hebert, D. J. Benjamin, J. Beauchamp, D. Cesarini, M. van der Loos, M. Johannesson, P. K. E. Magnusson, P. Lichtenstein, C. S. Atwood, J. Freese, T. S. Hauser, R. M. Hauser, N. Christakis, and D. Laibson. 2012. “Most Reported Genetic Associations With General Intelligence Are Probably False Positives.” Psychological Science 23(11):1314–23.

Consortium, The 1000 Genomes Project. 2015. “A Global Reference for Human Genetic Variation.” Nature 526(7571):68–74.

Davies, G., A. Tenesa, A. Payton, J. Yang, S. E. Harris, D. Liewald, X. Ke, S. Le Hellard, A. Christoforou, M. Luciano, K. McGhee, L. Lopez, A. J. Gow, J. Corley, P. Redmond, H. C. Fox, P. Haggarty, L. J. Whalley, G. McNeill, M. E. Goddard, T. Espeseth, A. J. Lundervold, I. Reinvang, A. Pickles, V. M. Steen, W. Ollier, D. J. Porteous, M. Horan, J. M. Starr, N. Pendleton, P. M. Visscher, and I. J. Deary. 2011. “Genome-wide association studies establish that human intelligence is highly heritable and polygenic.” Molecular Psychiatry 16(10):996–1005.

de Moor, M. H. M., P. T. Costa, A. Terracciano, R. F. Krueger, E. J. C. de Geus, T. Toshiko, Bwjh Penninx, T. Esko, P. A. F. Madden, J. Derringer, N. Amin, G. Willemsen, J. J. Hottenga, M. A. Distel, M. Uda, S. Sanna, P. Spinhoven, C. A. Hartman, P. Sullivan, A. Realo, J. Allik, A. C. Heath, M. L. Pergadia, A. Agrawal, P. Lin, R. Grucza, T. Nutile, M. Ciullo, D. Rujescu, I. Giegling, B. Konte, E. Widen, D. L. Cousminer, J. G. Eriksson, A. Palotie, L. Peltonen, M. Luciano, A. Tenesa, G. Davies, L. M. Lopez, N. K. Hansell, S. E. Medland, L. Ferrucci, D. Schlessinger, G. W. Montgomery, M. J. Wright, Y. S. Aulchenko, Acjw Janssens, B. A. Oostra, A. Metspalu, G. R. Abecasis, I. J. Deary, K. Raikkonen, L. J. Bierut, N. G. Martin, C. M. van Duijn, and D. I. Boomsma. 2012. “Meta-analysis of genome-wide association studies for personality.” Molecular Psychiatry 17(3):337–49.

Deary, I. J., S. Strand, P. Smith, and C. Fernandes. 2007. “Intelligence and educational achievement.” Intelligence 35(1):13–21.

DeGroot. 1948. “School was delayed for Dutch children.”

Della Bella, S., and M. Lucchini. 2015. “Education and BMI: a genetic informed analysis.” Quality & Quantity 49(6):2577–93.

Diggle, Peter J., Kung-Yee Liang, and Scott L. Zeger. 1994. Analysis of Longitudinal Data London: Oxford University Press.

Duncan, G. J., J. Brooksgunn, and P. K. Klebanov. 1994. “ECONOMIC DEPRIVATION AND EARLY-CHILDHOOD DEVELOPMENT.” Child Development 65(2):296–318.

Duncan, LE, H Shen, Bb Gelaye, K J Ressler, M W Feldman, RE Peterson, and B W Domingue. 2018. “Analysis of Polygenic Score Usage and Performance across Diverse Human Populations”. http://dx.doi.org/10.1101/398396.

Dunn, Lloyd M., and Douglas M. Dunn. 2007. Peabody Picture Vocabulary Test, Fourth Edition (PPVT™-4): Pearson Education.

Entwisle, D. R., and K. L. Alexander. 1992. “SUMMER SETBACK - RACE, POVERTY, SCHOOL COMPOSITION, AND MATHEMATICS ACHIEVEMENT IN THE 1ST 2 YEARS OF SCHOOL.” American Sociological Review 57(1):72–84.

Farkas, G., and K. Beron. 2004. “The detailed age trajectory of oral vocabulary knowledge: differences by class and race.” Social Science Research 33(3):464–97.

Farkas, G., P. England, K. Vicknair, and B. S. Kilbourne. 1997. “Cognitive skill, skill demands of jobs, and earnings among young European American, African American, and Mexican American workers.” Social Forces 75(3):913–38.

Farkas, G., and K. Vicknair. 1996. “Appropriate tests of racial wage discrimination require controls for cognitive skill: Comment on Cancio, Evans, and Maume.” American Sociological Review 61(4):557–60.

Fischer, Claude, Michael Hout, Martin Jankowski, Samuel Lucas, Ann Swidler, and Kim Voss. 1996. Inequality by Design: Cracking the Bell Curve Myth. Princeton, NJ: Princeton University Press.

Flynn, J. R. 1987. “MASSIVE IQ GAINS IN 14 NATIONS - WHAT IQ TESTS REALLY MEASURE.” Psychological Bulletin 101(2):171–91.

Flynn, J. R. 2009. “Requiem for nutrition as the cause of IQ gains: Raven’s gains in Britain 1938-2008.” Economics & Human Biology 7(1):18–27.

Flynn, James R. 2007. What Is Intelligence?: Beyond the Flynn Effect New York: Cambridge University Press

Frayling, T. M., N. J. Timpson, M. N. Weedon, E. Zeggini, R. M. Freathy, C. M. Lindgren, J. R. B. Perry, K. S. Elliott, H. Lango, N. W. Rayner, B. Shields, L. W. Harries, J. C. Barrett, S. Ellard, C. J. Groves, B. Knight, A. M. Patch, A. R. Ness, S. Ebrahim, D. A. Lawlor, S. M. Ring, Y. Ben-Shlomo, M. R. Jarvelin, U. Sovio, A. J. Bennett, D. Melzer, L. Ferrucci, R. J. F. Loos, I. Barroso, N. J. Wareham, F. Karpe, K. R. Owen, L. R. Cardon, M. Walker, G. A. Hitman, C. N. A. Palmer, A. S. F. Doney, A. D. Morris, G. D. Smith, A. T. Hattersley, and M. I. McCarthy. 2007. “A common variant in the FTO gene is associated with body mass index and predisposes to childhood and adult obesity.” Science 316(5826):889–94.

Furberg, H., Y. Kim, J. Dackor, E. Boerwinkle, N. Franceschini, D. Ardissino, L. Bernardinelli, P. M. Mannucci, F. Mauri, P. A. Merlini, D. Absher, T. L. Assimes, S. P. Fortmann, C. Iribarren, J. W. Knowles, T. Quertermous, L. Ferrucci, T. Tanaka, J. C. Bis, C. D. Furberg, T. Haritunians, B. McKnight, B. M. Psaty, K. D. Taylor, E. L. Thacker, P. Almgren, L. Groop, C. Ladenvall, M. Boehnke, A. U. Jackson, K. L. Mohlke, H. M. Stringham, J. Tuomilehto, E. J. Benjamin, S. J. Hwang, D. Levy, S. R. Preis, R. S. Vasan, J. Duan, P. V. Gejman, D. F. Levinson, A. R. Sanders, J. X. Shi, E. H. Lips, J. D. McKay, A. Agudo, L. Barzan, V. Bencko, S. Benhamou, X. Castellsague, C. Canova, D. I. Conway, E. Fabianova, L. Foretova, V. Janout, C. M. Healy, I. Holcatova, K. Kjaerheim, P. Lagiou, J. Lissowska, R. Lowry, T. V. Macfarlane, D. Mates, L. Richiardi, P. Rudnai, N. Szeszenia-Dabrowska, D. Zaridze, A. Znaor, M. Lathrop, P. Brennan, S. Bandinelli, T. M. Frayling, J. M. Guralnik, Y. Milaneschi, J. R. B. Perry, D. Altshuler, R. Elosua, S. Kathiresan, G. Lucas, O. Melander, C. J. O’Donnell, V. Salomaa, S. M. Schwartz, B. F. Voight, B. W. Penninx, J. H. Smit, N. Vogelzangs, D. I. Boomsma, E. J. C. de Geus, J. M. Vink, G. Willemsen, S. J. Chanock, F. Y. Gu, S. E. Hankinson, D. J. Hunter, A. Hofman, H. Tiemeier, A. G. Uitterlinden, C. M. van Duijn, S. Walter, et al. 2010. “Genome-wide meta-analyses identify multiple loci associated with smoking behavior.” Nature Genetics 42(5):441–U134.

Glazier, A. M., J. H. Nadeau, and T. J. Aitman. 2002. “Finding genes that underlie complex traits.” Science 298(5602):2345–49.

Gruer, L., C. L. Hart, and G. C. M. Watt. 2017. “After 50 years and 200 papers, what can the Midspan cohort studies tell us about our mortality?” Public Health 142:186–95.

Guo, G., and K. M. Harris. 2000. “The mechanisms mediating the effects of poverty on children's intellectual development.” Demography 37(4):431–47.

Guo, G., and E. Stearns. 2002. “The social influences on the realization of genetic potential for intellectual development.” Social Forces 80(3):881–910.

Harris, J. R. 1998. The nurture assumption: why children turn out the way they do. New York: Touchstone.

Hart, B., and R. T. Risley. 1995. Meaningful differences in the everyday experience of young American children Baltimore, MD: Paul H. Brookes.

Hauser, R. M. 2010. “Causes and Consequences of Cognitive Functioning Across the Life Course.” Educational Researcher 39(2):95–109.

Herrnstein, Richard J, and Charles Murray. 1994. The bell curve: Intelligence and class structure in American life. New York: Free Press.

Hindorff, L. A., V. L. Bonham, L. C. Brody, M. E. C. Ginoza, C. M. Hutter, T. A. Manolio, and E. D. Green. 2018. “Prioritizing diversity in human genomics research.” Nature Reviews Genetics 19(3):175-+.

Hinshaw, S. P. 1992. “EXTERNALIZING BEHAVIOR PROBLEMS AND ACADEMIC UNDERACHIEVEMENT IN CHILDHOOD AND ADOLESCENCE - CAUSAL RELATIONSHIPS AND UNDERLYING MECHANISMS.” Psychological Bulletin 111(1):127–55.

Ickovics, J. R., A. Carroll-Scott, S. M. Peters, M. Schwartz, K. Gilstad-Hayden, and C. McCaslin. 2014. “Health and Academic Achievement: Cumulative Effects of Health Assets on Standardized Test Scores Among Urban Youth in the United States.” Journal of School Health 84(1):40–48.

Jencks, Christopher, Susan Bartlett Bartlett, Mary Corcoran, James Crouse, David Eaglesfield, Gregory Jackson, Kent McClelland McClelland, Peter Mueser, Michael Olneck, Joseph Schwarz, Sherry Ward, and Jill Williams. 1979. Who Gets Ahead? The Determinants of Economic Success in America. New York: Basic Books.

Jensen, A. R. 1969. “HOW MUCH CAN WE BOOST IQ AND SCHOLASTIC ACHIEVEMENT.” Harvard Educational Review 39(1):1–123.

Jensen, Arthur R. 1997. “The puzzle of nongenetic variance.” Intelligence, heredity, and environment:42–88.

Kerckhoff, A. C., S. W. Raudenbush, and E. Glennie. 2001. “Education, cognitive skill, and labor force outcomes.” Sociology of Education 74(1):1–24.

Kessler, R. C., C. L. Foster, W. B. Saunders, and P. E. Stang. 1995. “SOCIAL-CONSEQUENCES OF PSYCHIATRIC-DISORDERS .1. EDUCATIONAL-ATTAINMENT.” American Journal of Psychiatry 152(7):1026–32.

Kong, A., G. Thorleifsson, M. L. Frigge, B. J. Vilhjalmsson, A. I. Young, T. E. Thorgeirsson, S. Benonisdottir, A. Oddsson, B. V. Halldorsson, G. Masson, D. F. Gudbjartsson, A. Helgason, G. Bjornsdottir, U. Thorsteinsdottir, and K. Stefansson. 2018. “The nature of nurture: Effects of parental genotypes.” Science 359(6374):424–28.

Lee, J. J., R. Wedow, A. Okbay, E. Kong, O. Maghzian, M. Zacher, T. A. Nguyen-Viet, P. Bowers, J. Sidorenko, R. K. Linner, M. A. Fontana, T. Kundu, C. Lee, H. Li, R. X. Li, R. Royer, P. N. Timshel, R. K. Walters, E. A. Willoughby, L. Yengo, M. Alver, Y. C. Bao, D. W. Clark, F. R. Day, N. A. Furlotte, P. K. Joshi, K. E. Kemper, A. Kleinman, C. Langenberg, R. Magi, J. W. Trampush, S. S. Verma, Y. Wu, M. Lam, J. H. Zhao, Z. L. Zheng, J. D. Boardman, H. Campbell, J. Freese, K. M. Harris, C. Hayward, P. Herd, M. Kumari, T. Lencz, J. A. Luan, A. K. Malhotra, A. Metspalu, L. Milani, K. K. Ong, J. R. B. Perry, D. J. Porteous, M. D. Ritchie, M. C. Smart, B. H. Smith, J. Y. Tung, N. J. Wareham, J. F. Wilson, J. P. Beauchamp, D. C. Conley, T. Esko, S. F. Lehrer, P. K. E. Magnusson, S. Oskarsson, T. H. Pers, M. R. Robinson, K. Thom, C. Watson, C. F. Chabris, M. N. Meyer, D. I. Laibson, J. Yang, M. Johannesson, P. D. Koellinger, P. Turley, P. M. Visscher, D. J. Benjamin, D. Cesarini, Team Me Res, Cogent Cognitive Genomics Consorti, and Consortiu Social Sci Genetic Assoc. 2018. “Gene discovery and polygenic prediction from a genome-wide association study of educational attainment in 1.1 million individuals.” Nature Genetics 50(8):1112-+.

Lemann, N. 1999. The Big Test: The Secret History of the American Meritocracy New York: Straus and Giroux.

Levitt, S. D., and S. J. Dubner. 2006. Freakonomics: A rogue economist explores the hidden side of everything. New York: William Morrow.

Link, B. G., and J. C. Phelan. 1996. “Understanding sociodemographic differences in health - The role of fundamental social causes.” American Journal of Public Health 86(4):471–73.

Locke, A. E., B. Kahali, S. I. Berndt, A. E. Justice, T. H. Pers, R. Felix, C. Powell, S. Vedantam, M. L. Buchkovich, J. Yang, D. C. Croteau-Chonka, T. Esko, T. Fall, T. Ferreira, S. Gustafsson, Z. Kutalik, J. A. Luan, R. Magi, J. C. Randall, T. W. Winkler, A. R. Wood, T. Workalemahu, J. D. Faul, J. A. Smith, J. H. Zhao, W. Zhao, J. Chen, R. Fehrmann, A. K. Hedman, J. Karjalainen, E. M. Schmidt, D. Absher, N. Amin, D. Anderson, M. Beekman, J. L. Bolton, L. Bragg-Gresham, S. Buyske, A. Demirkan, G. H. Deng, G. B. Ehret, B. Feenstra, M. F. Feitosa, K. Fischer, A. Goel, J. Gong, A. U. Jackson, S. Kanoni, M. E. Kleber, K. Kristiansson, U. Lim, V. Lotay, M. Mangino, I. M. Leach, C. Medina-Gomez, S. E. Medland, M. A. Nalls, C. D. Palmer, D. Pasko, S. Pechlivanis, M. J. Peters, I. Prokopenko, D. Shungin, A. Stancakova, R. J. Strawbridge, Y. J. Sung, T. Tanaka, A. Teumer, S. Trompet, S. W. van der Laan, J. van Settee, J. V. Van Vliet-Ostaptchouk, Z. M. Wang, L. Yengo, W. H. Zhang, A. Isaacs, E. Albrecht, J. Arnlov, G. M. Arscott, A. P. Attwood, S. Bandinelli, A. Barrett, I. N. Bas, C. Bellis, A. J. Bennett, C. Berne, R. Blagieva, M. Bluher, S. Bohringer, L. L. Bonnycastle, Y. Bottcher, H. A. Boyd, M. Bruinenberg, I. H. Caspersen, Y. D. Chen, R. Clarke, E. W. Daw, A. J. M. de Craen, G. Delgado, M. Dimitriou, et al. 2015. “Genetic studies of body mass index yield new insights for obesity biology.” Nature 518(7538):197–U401.

Locurto, C. 1990. “THE MALLEABILITY OF IQ AS JUDGED FROM ADOPTION STUDIES.” Intelligence 14(3):275–92.

Lynn, R., and I. E. Gordon. 1961. “THE RELATION OF NEUROTICISM AND EXTRAVERSION TO INTELLIGENCE AND EDUCATIONAL-ATTAINMENT.” British Journal of Educational Psychology 31(2):194–203.

MacArthur, Jacqueline, Emily Bowler, Maria Cerezo, Laurent Gil, Peggy Hall, Emma Hastings, Heather Junkins, Aoife McMahon, Annalisa Milano, Joannella Morales, Zoe May Pendlington, Danielle Welter, Tony Burdett, Lucia Hindorff, Paul Flicek, Fiona Cunningham, and Helen Parkinson. 2017. “The New NHGRI-EBI Catalog of Published Genome-Wide Association Studies (Gwas Catalog).” Nucleic Acids Research 45:D896–D901.

Malinauskiene, O., R. Vosylis, R. Zukauskiene, and B. V. Elsevier Science. 2011. “Longitudinal examination of relationships between problem behaviors and academic achievement in young adolescents.” Pp. 3415–21 in 3rd World Conference on Educational Sciences.

Marouli, E., M. Graff, C. Medina-Gomez, K. S. Lo, A. R. Wood, T. R. Kjaer, R. S. Fine, Y. C. Lu, C. Schurmann, H. M. Highland, S. Rueger, G. Thorleifsson, A. E. Justice, D. Lamparter, K. E. Stirrups, V. Turcot, K. L. Young, T. W. Winkler, T. Esko, T. Karaderi, A. E. Locke, N. G. D. Masca, M. C. Y. Ng, P. Mudgal, M. A. Rivas, S. Vedantam, A. Mahajan, X. Q. Guo, G. Abecasis, K. K. Aben, L. S. Adair, D. S. Alam, E. Albrecht, K. H. Allin, M. Allison, P. Amouyel, E. V. Appel, D. Arveiler, F. W. Asselbergs, P. L. Auer, B. Balkau, B. Banas, L. E. Bang, M. Benn, S. Bergmann, L. F. Bielak, M. Bluher, H. Boeing, E. Boerwinkle, C. A. Boger, L. L. Bonnycastle, J. Bork-Jensen, M. L. Bots, E. P. Bottinger, D. W. Bowden, I. Brandslund, G. Breen, M. H. Brilliant, L. Broer, A. A. Burt, A. S. Butterworth, D. J. Carey, M. J. Caulfield, J. C. Chambers, D. I. Chasman, Y. D. I. Chen, R. Chowdhury, C. Christensen, A. Y. Chu, M. Cocca, F. S. Collins, J. P. Cook, J. Corley, J. C. Galbany, A. J. Cox, G. Cuellar-Partida, J. Danesh, G. Davies, P. I. W. de Bakker, G. J. de Borst, S. de Denus, M. C. H. de Groot, R. de Mutsert, I. J. Deary, G. Dedoussis, E. W. Demerath, A. I. den Hollander, J. G. Dennis, E. Di Angelantonio, F. Drenos, M. M. Du, A. M. Dunning, D. F. Easton, T. Ebeling, T. L. Edwards, P. T. Ellinor, P. Elliott, E. Evangelou, A. E. Farmaki, J. D. Faul, et al. 2017. “Rare and low-frequency coding variants alter human adult height.” Nature 542(7640):186–90.

McEwen, B. S. 2000. “The neurobiology of stress: from serendipity to clinical relevance.” Brain Research 886(1-2):172–89.

McKenzie, J., M. Taghavi-Khonsary, and G. Tindell. 2000. “Neuroticism and academic achievement: the Furneaux Factor as a measure of academic rigour.” Personality and Individual Differences 29(1):3–11.

McLeod, J. D., and K. Kaiser. 2004. “Childhood emotional and behavioral problems and educational attainment.” American Sociological Review 69(5):636–58.

McLeod, J. D., and M. J. Shanahan. 1993. “POVERTY, PARENTING, AND CHILDRENS MENTAL-HEALTH.” American Sociological Review 58(3):351–66.

Nisbett, Richard E. 2009. Intelligence and how to get it. New York: W.W. Norton & Company, Inc.

Okbay, A., J. P. Beauchamp, M. A. Fontana, J. J. Lee, T. H. Pers, C. A. Rietveld, P. Turley, G. B. Chen, V. Emilsson, S. F. W. Meddens, S. Oskarsson, J. K. Pickrell, K. Thom, P. Timshel, R. de Vlaming, A. Abdellaoui, T. S. Ahluwalia, J. Bacelis, C. Baumbach, G. Bjornsdottir, J. H. Brandsma, M. P. Concas, J. Derringer, N. A. Furlotte, T. E. Galesloot, G. Girotto, R. Gupta, L. M. Hall, S. E. Harris, E. Hofer, M. Horikoshi, J. E. Huffman, K. Kaasik, I. P. Kalafati, R. Karlsson, A. Kong, J. Lahti, S. J. van der Lee, C. de Leeuw, P. A. Lind, K. O. Lindgren, T. Liu, M. Mangino, J. Marten, E. Mihailov, M. B. Miller, P. J. van der Most, C. Oldmeadow, A. Payton, N. Pervjakova, W. J. Peyrot, Y. Qian, O. Raitakari, R. Rueedi, E. Salvi, B. Schmidt, K. E. Schraut, J. X. Shi, A. V. Smith, R. A. Poot, B. St Pourcain, A. Teumer, G. Thorleifsson, N. Verweij, D. Vuckovic, J. Wellmann, H. J. Westra, J. Y. Yang, W. Zhao, Z. H. Zhu, B. Z. Alizadeh, N. Amin, A. Bakshi, S. E. Baumeister, G. Biino, K. Bonnelykke, P. A. Boyle, H. Campbell, F. P. Cappuccio, G. Davies, J. E. De Neve, P. Deloukas, I. Demuth, J. Ding, P. Eibich, L. Eisele, N. Eklund, D. M. Evans, J. D. Faul, M. F. Feitosa, A. J. Forstner, I. Gandin, B. Gunnarsson, B. V. Halldorsson, T. B. Harris, A. C. Heath, L. J. Hocking, E. G. Holliday, G. Homuth, M. A. Horan, et al. 2016. “Genome-wide association study identifies 74 loci associated with educational attainment.” Nature 533(7604):539-+.

Phillips, J. B., and N. S. Endler. 1982. “ACADEMIC EXAMINATIONS AND ANXIETY - THE INTERACTION-MODEL EMPIRICALLY TESTED.” Journal of Research in Personality 16(3):303–18.

Pietschnig, J., L. Penke, J. M. Wicherts, M. Zeiler, and M. Voracek. 2015. “Meta-analysis of associations between human brain volume and intelligence differences: How strong are they and what do they mean?" Neuroscience and Biobehavioral Reviews 57:411–32.

Pinker, S. 2002. The blank slate: The modern denial of human nature. New York: Viking.

Plomin, R., and J. C. Loehlin. 1989. “DIRECT AND INDIRECT IQ HERITABILITY ESTIMATES - A PUZZLE.” Behavior Genetics 19(3):331–42.

Plomin, Robert. 2018. “Genetics ‘very important’ in shaping lives.” BBC World News HARDtalk.

Price, A. L., N. J. Patterson, R. M. Plenge, M. E. Weinblatt, N. A. Shadick, and D. Reich. 2006. “Principal components analysis corrects for stratification in genome-wide association studies.” Nature Genetics 38:904–09.

Raudenbush, S. W, and A. S. Bryk. 2002. Hierarchical Linear Models: Applications and Data Analysis Methods Thousand Oaks, CA: Sage.

Rietveld, C. A., S. E. Medland, J. Derringer, J. Yang, T. Esko, N. W. Martin, H. J. Westra, K. Shakhbazov, A. Abdellaoui, A. Agrawal, E. Albrecht, B. Z. Alizadeh, N. Amin, J. Bamard, S. E. Baumeister, K. S. Benke, L. F. Bielak, J. A. Boatman, P. A. Boyle, G. Davies, C. De Leeuw, N. Eklund, D. S. Evans, R. Ferhmann, K. Fischer, C. Gieger, H. K. Gjessing, S. Hagg, J. R. Harris, C. Hayward, C. Holzapfel, C. A. Ibrahim-Verbaas, E. Ingelsson, B. Jacobsson, P. K. Joshi, A. Jugessur, M. Kaakinen, S. Kanoni, J. Karjalainen, I. Kolcic, K. Kristiansson, Z. Kutalik, J. Lahti, S. H. Lee, P. Lin, P. A. Lind, Y. M. Liu, K. Lohman, M. Loitfelder, G. McMahon, P. M. Vidal, O. Meirelles, L. Milani, R. Myhre, M. L. Nuotio, C. J. Oldmeadow, K. E. Petrovic, W. J. Peyrot, O. Polasek, L. Quaye, E. Reinmaa, J. P. Rice, T. S. Rizzi, H. Schmidt, R. Schmidt, A. V. Smith, J. A. Smith, T. Tanaka, A. Terracciano, Mjhm van der Loos, V. Vitart, H. Volzke, J. Wellmann, L. Yu, W. Zhao, J. Allik, J. R. Attia, S. Bandinelli, F. Bastardot, J. Beauchamp, D. A. Bennett, K. Berger, L. J. Bierut, D. I. Boomsma, U. Bultmann, H. Campbell, C. F. Chabris, L. Cherkas, M. K. Chung, F. Cucca, M. de Andrade, P. L. De Jager, J. E. De Neve, I. J. Deary, G. V. Dedoussis, P. Deloukas, M. Dimitriou, G. Eiriksdottir, M. F. Elderson, J. G. Eriksson, et al. 2013. “GWAS of 126,559 Individuals Identifies Genetic Variants Associated with Educational Attainment.” Science 340(6139):1467–71.

Rosenbaum, J.. 2001. Beyond College for All: Career Paths for the Forgotten Half. New York: Russell Sage Found.

Roskam, A. J. R., A. E. Kunst, H. Van Oyen, S. Demarest, J. Klumbiene, E. Regidor, U. Helmert, F. Jusot, D. Dzurova, and J. P. Mackenbach. 2010. “Comparative appraisal of educational inequalities in overweight and obesity among adults in 19 European countries.” International Journal of Epidemiology 39(2):392–404.

Savage, J. E., P. R. Jansen, S. Stringer, K. Watanabe, J. Bryois, C. A. de Leeuw, M. Nagel, S. Awasthi, P. B. Barr, J. R. I. Coleman, K. L. Grasby, A. R. Hammerschlag, J. A. Kaminski, R. Karlsson, E. Krapohl, M. Lam, M. Nygaard, C. A. Reynolds, J. W. Trampush, H. Young, D. Zabaneh, S. Hagg, N. K. Hansell, I. K. Karlsson, S. Linnarsson, G. W. Montgomery, A. B. Munoz-Manchado, E. B. Quinlan, G. Schumann, N. G. Skene, B. T. Webb, T. White, D. E. Arking, D. Avramopoulos, R. M. Bilder, P. Bitsios, K. E. Burdick, T. D. Cannon, O. Chiba-Falek, A. Christoforou, E. T. Cirulli, E. Congdon, A. Corvin, G. Davies, I. J. Deary, P. DeRosse, D. Dickinson, S. Djurovic, G. Donohoe, E. D. Conley, J. G. Eriksson, T. Espeseth, N. A. Freimer, S. Giakoumaki, I. Giegling, M. Gill, D. C. Glahn, A. R. Hariri, A. Hatzimanolis, M. C. Keller, E. Knowles, D. Koltai, B. Konte, J. Lahti, S. Le Hellard, T. Lencz, D. C. Liewald, E. London, A. J. Lundervold, A. K. Malhotra, I. Melle, D. Morris, A. C. Need, W. Ollier, A. Palotie, A. Payton, N. Pendleton, R. A. Poldrack, K. Raikkonen, I. Reinvang, P. Roussos, D. Rujescu, F. W. Sabb, M. A. Scult, O. B. Smeland, N. Smyrnis, J. M. Starr, V. M. Steen, N. C. Stefanis, R. E. Straub, K. Sundet, H. Tiemeier, A. N. Voineskos, D. R. Weinberger, E. Widen, J. Yu, G. Abecasis, O. A. Andreassen, G. Breen, L. Christiansen, et al. 2018. “Genome-wide association meta-analysis in 269,867 individuals identifies new genetic and functional links to intelligence.” Nature Genetics 50(7):912-+.

Scullin, M. H., E. Peters, W. M. Williams, and S. J. Ceci. 2000. “The role of IQ and education in predicting later labor market outcomes - Implications for affirmative action.” Psychology Public Policy and Law 6(1):63–89.

Searle, S. R. 1971. Linear Models. New York: Wiley and Sons.

Searle, S.R., G Casella, and C McCulloch. 1992. Variance Components New York: Wiley & Sons.

Sniekers, S., S. Stringer, K. Watanabe, P. R. Jansen, J. R. I. Coleman, E. Krapohl, E. Taskesen, A. R. Hammerschlag, A. Okbay, D. Zabaneh, N. Amin, G. Breen, D. Cesarini, C. F. Chabris, W. G. Iacono, M. A. Ikram, M. Johannesson, P. Koellinger, J. J. Lee, P. K. E. Magnusson, M. McGue, M. B. Miller, W. E. R. Ollier, A. Payton, N. Pendleton, R. Plomin, C. A. Rietveld, H. Tiemeier, C. M. van Duijn, and D. Posthuma. 2017. “Genome-wide association meta-analysis of 78,308 individuals identifies new loci and genes influencing human intelligence.” Nature Genetics 49(7):1107-+.

Speliotes, Elizabeth K., Cristen J. Willer, Sonja I. Berndt, Keri L. Monda, Gudmar Thorleifsson, Anne U. Jackson, Hana Lango Allen, Cecilia M. Lindgren, Jian'an Luan, Reedik Maegi, Joshua C. Randall, Sailaja Vedantam, Thomas W. Winkler, Lu Qi, Tsegaselassie Workalemahu, Iris M. Heid, Valgerdur Steinthorsdottir, Heather M. Stringham, Michael N. Weedon, Eleanor Wheeler, Andrew R. Wood, Teresa Ferreira, Robert J. Weyant, Ayellet V. Segre, Karol Estrada, Liming Liang, James Nemesh, Ju-Hyun Park, Stefan Gustafsson, Tuomas O. Kilpelaenen, Jian Yang, Nabila Bouatia-Naji, Tonu Esko, Mary F. Feitosa, Zoltan Kutalik, Massimo Mangino, Soumya Raychaudhuri, Andre Scherag, Albert Vernon Smith, Ryan Welch, Jing Hua Zhao, Katja K. Aben, Devin M. Absher, Najaf Amin, Anna L. Dixon, Eva Fisher, Nicole L. Glazer, Michael E. Goddard, Nancy L. Heard-Costa, Volker Hoesel, Jouke-Jan Hottenga, Asa Johansson, Toby Johnson, Shamika Ketkar, Claudia Lamina, Shengxu Li, Miriam F. Moffatt, Richard H. Myers, Narisu Narisu, John R. B. Perry, Marjolein J. Peters, Michael Preuss, Samuli Ripatti, Fernando Rivadeneira, Camilla Sandholt, Laura J. Scott, Nicholas J. Timpson, Jonathan P. Tyrer, Sophie van Wingerden, Richard M. Watanabe, Charles C. White, Fredrik Wiklund, Christina Barlassina, Daniel I. Chasman, Matthew N. Cooper, John-Olov Jansson, Robert W. Lawrence, Niina Pellikka, Inga Prokopenko, Jianxin Shi, Elisabeth Thiering, Helene Alavere, Maria T. S. Alibrandi, Peter Almgren, Alice M. Arnold, Thor Aspelund, Larry D. Atwood, Beverley Balkau, Anthony J. Balmforth, Amanda J. Bennett, Yoav Ben-Shlomo, Richard N. Bergman, Sven Bergmann, Heike Biebermann, Alexandra I. F. Blakemore, Tanja Boes, Lori L. Bonnycastle, Stefan R. Bornstein, Morris J. Brown, Thomas A. Buchanan, et al. 2010. “Association analyses of 249,796 individuals reveal 18 new loci associated with body mass index.” Nature Genetics 42(11):937–U53.

Spilerman, S., and T. Lunde. 1991. “FEATURES OF EDUCATIONAL-ATTAINMENT AND JOB PROMOTION PROSPECTS.” American Journal of Sociology 97(3):689–720.

Taal, H Rob, Beate St Pourcain, Elisabeth Thiering, Shikta Das, Dennis O Mook-Kanamori, Nicole M Warrington, Marika Kaakinen, Eskil Kreiner-Møller, Jonathan P Bradfield, Rachel M Freathy, Frank Geller, Mònica Guxens, Diana L Cousminer, Marjan Kerkhof, Nicholas J Timpson, M Arfan Ikram, Lawrence J Beilin, Klaus Bønnelykke, Jessica L Buxton, Pimphen Charoen, Bo Lund Krogsgaard Chawes, Johan Eriksson, David M Evans, Albert Hofman, John P Kemp, Cecilia E Kim, Norman Klopp, Jari Lahti, Stephen J Lye, George McMahon, Frank D Mentch, Martina Müller-Nurasyid, Paul F O'Reilly, Inga Prokopenko, Fernando Rivadeneira, Eric A P Steegers, Jordi Sunyer, Carla Tiesler, Hanieh Yaghootkar, The Cohorts for Heart Consortium, Aging Research in Genetic Epidemiology, M Arfan Ikram, Myriam Fornage, Albert V Smith, Sudha Seshadri, Reinhold Schmidt, Stéphanie Debette, Henri A Vrooman, Sigurdur Sigurdsson, Stefan Ropele, Laura H Coker, W T Longstreth Jr, Wiro J Niessen, Anita L DeStefano, Alexa Beiser, Alex P Zijdenbos, Maksim Struchalin, Clifford R Jack Jr, Mike A Nalls, Rhoda Au, Albert Hofman, Haukur Gudnason, Aad van der Lugt, Tamara B Harris, William M Meeks, Meike W Vernooij, Mark A van Buchem, Diane Catellier, Vilmundur Gudnason, B Gwen Windham, Philip A Wolf, Cornelia M van Duijn, Thomas H Mosley Jr, Helena Schmidt, Lenore J Launer, Monique M B Breteler, Charles DeCarli, Monique M B Breteler, Stéphanie Debette, Myriam Fornage, Vilmundur Gudnason, Lenore J Launer, Aad van der Lugt, Thomas H Mosley Jr, Sudha Seshadri, Albert V Smith, Meike W Vernooij, Early Genetics Consortium, Lifecourse Epidemiology, Wei Ang, Toos van Beijsterveldt, Nienke Bergen, Kelly Benke, Diane Berry, Jonathan P Bradfield, Pimphen Charoen, Lachlan Coin, Diana L Cousminer, Shikta Das, Paul Elliott, et al. 2012. “Common variants at 12q15 and 12q24 are associated with infant head circumference.” Nature Genetics 44(5):532–38.

Taubman, P, and T. Wales. 1974. Higher Education and Earnings. New York: McGraw-Hill.

Taubman, Paul, and Terence Wales. 1972. Mental Ability and Higher Educational Attainment in the 20th Century: NBER Books, National Bureau of Economic Research, Inc.

Thornberry, T. P., and M. D. Krohn. 2000. “The self-report method for measuring delinquency and crime.” Pp. 33–83 in Criminal Justice 2000. Washington, D.C.: National Institute of Justice.

Turkheimer, E., A. Haley, M. Waldron, B. D'Onofrio, and Gottesman, II. 2003. “Socioeconomic status modifies heritability of IQ in young children.” Psychological Science 14(6):623–28.

Velden, M. 1997. “The heritability of intelligence: Neither known nor unknown.” American Psychologist 52(1):72–73.

Winship, Christopher, and Sanders Korenman. 1997. “Does staying in school make you smarter? The effect of education on IQ in The Bell Curve” Pp. 215–34 in Intelligence and Success: Is it all in the Genes? Scientists Respond to The Bell Curve., edited by Bernie Devlin, Stephen E. Fienberg, Daniel Resnick, and Kathryn Reder: Springer-Verlag.

Wodtke, Geoffrey T., David J. Harding, and Felix Elwert. 2011. “Neighborhood Effects in Temporal Perspective: The Impact of Long-Term Exposure to Concentrated Disadvantage on High School Graduation.” American Sociological Review 76(5):713–36.

Wood, Andrew R., Tonu Esko, Jian Yang, Sailaja Vedantam, Tune H. Pers, Stefan Gustafsson, Audrey Y. Chun, Karol Estrada, Jian'an Luan, Zoltan Kutalik, Najaf Amin, Martin L. Buchkovich, Damien C. Croteau-Chonka, Felix R. Day, Yanan Duan, Tove Fall, Rudolf Fehrmann, Teresa Ferreira, Anne U. Jackson, Juha Karjalainen, Ken Sin Lo, Adam E. Locke, Reedik Maegi, Evelin Mihailov, Eleonora Porcu, Joshua C. Randall, Andre Scherag, Anna A. E. Vinkhuyzen, Harm-Jan Westra, Thomas W. Winkler, Tsegaselassie Workalemahu, Jing Hua Zhao, Devin Absher, Eva Albrecht, Denise Anderson, Jeffrey Baron, Marian Beekman, Ayse Demirkan, Georg B. Ehret, Bjarke Feenstra, Mary F. Feitosa, Krista Fischer, Ross M. Fraser, Anuj Goel, Jian Gong, Anne E. Justice, Stavroula Kanoni, Marcus E. Kleber, Kati Kristiansson, Unhee Lim, Vaneet Lotay, Julian C. Lui, Massimo Mangino, Irene Mateo Leach, Carolina Medina-Gomez, Michael A. Nalls, Dale R. Nyholt, Cameron D. Palmer, Dorota Pasko, Sonali Pechlivanis, Inga Prokopenko, Janina S. Ried, Stephan Ripke, Dmitry Shungin, Alena Stancakova, Rona J. Strawbridge, Yun Ju Sung, Toshiko Tanaka, Alexander Teumer, Stella Trompet, Sander W. van der Laan, Jessica van Setten, Jana V. Van Vliet-Ostaptchouk, Zhaoming Wang, Loic Yengo, Weihua Zhang, Uzma Afzal, Johan Arnloev, Gillian M. Arscott, Stefania Bandinelli, Amy Barrett, Claire Bellis, Amanda J. Bennett, Christian Berne, Matthias Blueher, Jennifer L. Bolton, Yvonne Boettcher, Heather A. Boyd, Marcel Bruinenberg, Brendan M. Buckley, Steven Buyske, Ida H. Caspersen, Peter S. Chines, Robert Clarke, Simone Claudi-Boehm, Matthew Cooper, E. Warwick Daw, Pim A. De Jong, Joris Deelen, Graciela Delgado, et al. 2014. “Defining the role of common variation in the genomic and biological architecture of adult human height.” Nature Genetics 46(11):1173–86.

Wraw, C., I. J. Deary, C. R. Gale, and G. Der. 2015. “Intelligence in youth and health at age 50.” Intelligence 53:23–32.

Yang, Yang Claire, Karen Gerken, Kristen Schorpp, Courtney Boen, and Kathleen Mullan Harris. 2017. “Early-Life Socioeconomic Status and Adult Physiological Functioning: A Life Course Examination of Biosocial Mechanisms.” Biodemography and Social Biology 63(2):87–103.

Zeggini, E., M. N. Weedon, C. M. Lindgren, T. M. Frayling, K. S. Elliott, H. Lango, N. J. Timpson, J. R. B. Perry, N. W. Rayner, R. M. Freathy, J. C. Barrett, B. Shields, A. P. Morris, S. Ellard, C. J. Groves, L. W. Harries, J. L. Marchini, K. R. Owen, B. Knight, L. R. Cardon, M. Walker, G. A. Hitman, A. D. Morris, A. S. F. Doney, M. I. McCarthy, and A. T. Hattersley. 2007. “Replication of genome-wide association signals in UK samples reveals risk loci for type 2 diabetes.” Science 316(5829):1336–41.

